# Unsaturated fatty acid synthesis is associated with poor prognosis and differentially regulated by *MYCN* and tumor suppressor microRNAs in neuroblastoma

**DOI:** 10.1101/2023.04.20.537692

**Authors:** Dennis A. Sheeter, Secilia Garza, Hui Gyu Park, Niharika R. Badi, Erika C. Espinosa, Kumar Kothapalli, J. Thomas Brenna, John T. Powers

## Abstract

*MYCN* amplification and disruption of tumor suppressor microRNA (TSmiR) function are central drivers of poor outcomes in neuroblastoma (NB). MYC, MYCN, and TSmiRs regulate glucose metabolism; however, their role in unsaturated fatty acid synthesis (UFAS) remains poorly understood. Here we show that *de novo* and UFAS pathway genes *FASN*, *ELOVL6*, *SCD*, *FADS2*, and *FADS1* are upregulated in high-risk NB and are associated with poor prognosis. RNA-Seq analysis of eight human NB cell lines revealed parallel UFAS gene expression patterns. Consistent with this, we found that NB-related TSmiRs were predicted to extensively target these genes. In addition, we observed that both MYC and MYCN upregulated UFAS pathway genes while suppressing TSmiR host gene expression, thereby creating a possible UFAS regulatory network between *MYCN* and TSmiRs in NB. Furthermore, NB cells are high in omega 9 (ω9) unsaturated fatty acids that can be synthesized *de novo* and low in both ω6 and ω3, providing a plausible means for NB to limit cell-autonomous immune stimulation and reactive oxygen species (ROS)-driven apoptosis from ω6 and ω3 unsaturated fatty acid derivatives, respectively. We propose a model in which the UFAS pathway, through novel regulation by *MYCN* and TSmiRs, plays a key role in neuroblastoma pathology with implications for other *MYC*-driven cancers.

## INTRODUCTION

Neuroblastoma (NB) is a highly metastatic pediatric cancer derived from the neural crest-sympathoadrenal lineage that normally contributes to the sympathetic ganglia, chromaffin cells of the adrenal medulla, and paraganglia (1). *MYCN* is the defining oncogene of NB, where it was first discovered, receiving its *MYC-“N”* designation from NB itself (2). *MYCN* gene amplification is a hallmark of high-risk and poor prognosis, where overall patient survival is below 50% (3–5). The MYCN and MYC transcription factors have highly overlapping function (2,6). MYC overexpression drives metabolic conversion to favor the rapid generation of ATP via anaerobic glycolysis over oxidative phosphorylation, a shift known as the Warburg effect (**Figure 1a**). High lactate dehydrogenase (LDH), known to be transcriptionally regulated by MYC, is a hallmark of this effect, shunting pyruvate use away from mitochondrial oxidative phosphorylation towards production and extracellular export of lactate (7,8). The remaining pyruvate enters the mitochondria where it is converted to citrate and exported. Increased mitochondrial citrate efflux raises cytoplasmic citrate levels, and ACLY catalyzes the release of acetyl-CoA, the fundamental building block for cytoplasmic *de novo* fatty acid synthesis (**Figure 1b**) (8). MYC has been implicated in fatty acid (FA) synthesis (FAS) via glycolysis-driven enhancement of acetyl-CoA production and increased expression of *de novo* FAS genes *ACLY*, *ACACA*, *FASN,* and *SCD* (**Figure 1b**) (9). Similar to *MYC*, *MYCN* is associated with both enhanced glycolysis and metabolic reprogramming, and has also been implicated in fatty acid uptake and *de novo* synthesis in NB (10–12).

**Figure 1.**
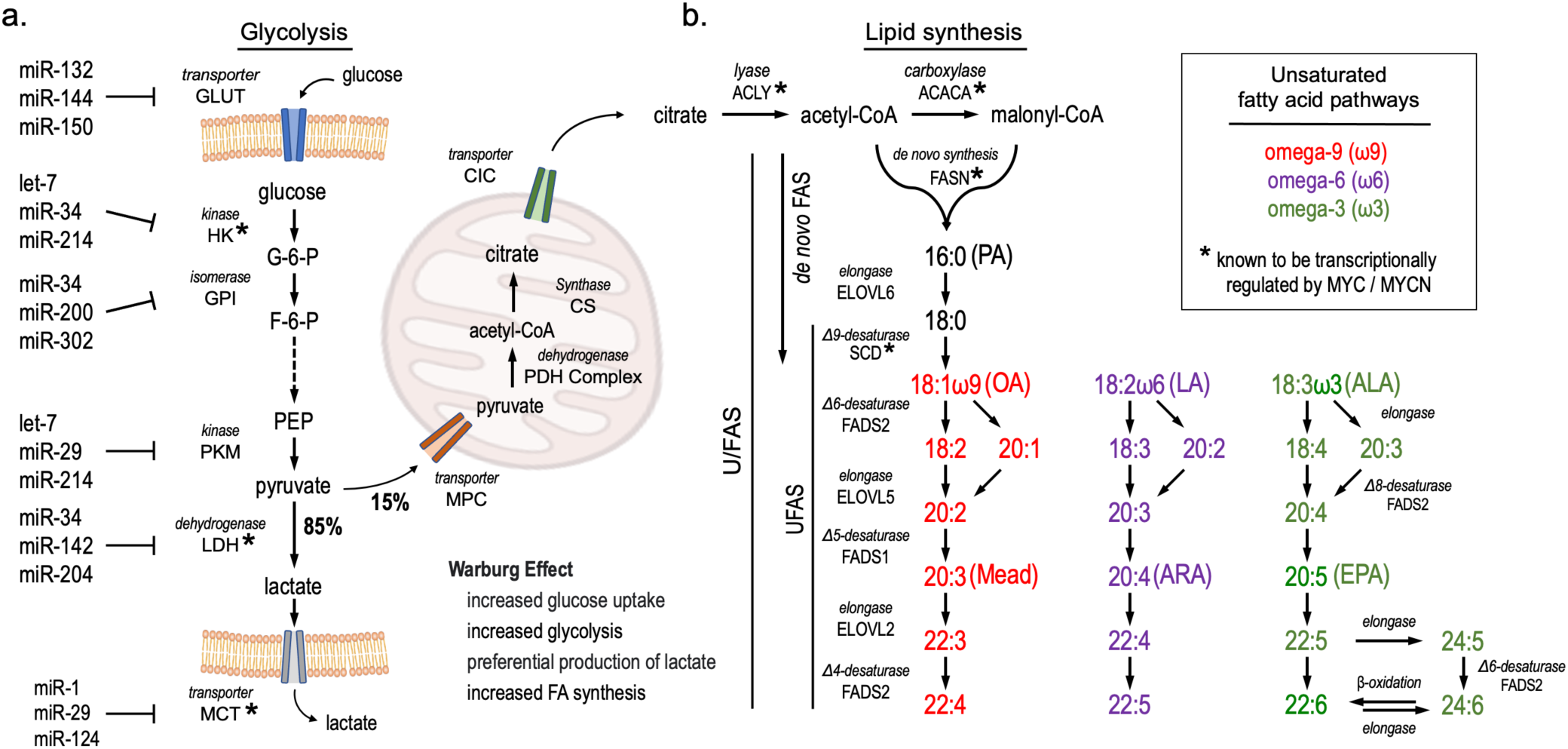
Glycolysis and fatty acid synthesis pathways. **a.** Schematic of glycolysis genes and known microRNA regulators. Enzymes regulated by MYC are denoted with an asterisk (*). **b.** Schematic of fatty acid synthesis. FASN catalyzes *de novo* FA synthesis from acetyl-CoA and malonyl-CoA subunits up to sixteen unsaturated carbon chain palmitic acid (16:0, PA), which then enters elongation and desaturation pathways to yield oleic acid (18:1ω9, OA) which can continue along the *de novo* pathway to Mead acid (20:3ω9) and its elongation product, DTrA (22:3ω9). Essential ω6 linoleic acid (18:2ω6, LA) and alpha linolenic acid (18:3ω3, ALA) are elongated and desaturated by the same set of enzymes.

In addition to their role as energy substrates, fatty acids are the main structural components of all cellular membranes. Specific non-esterified free fatty acids are key to intracellular and paracrine cell signaling, as well as when acylated to phosphatidylinositol and diacylglycerol. Many paracrine signals are immune-stimulating, including prostaglandins, 20-HETE, and leukotrienes (13). The relative proportions of the fatty acyl chains making up membrane phospholipids has a major influence on the signaling molecules produced on an ‘as needed’ basis in response to stimulation (14). This constraint places extraordinary importance on the regulation of fatty acid-derived signaling molecule production, which has implications for the uptake, synthesis, downstream processing, and relative proportion of individual fatty acids within cell membranes. With respect to cancer, fatty acid profiles can be an effect or cause; transformation to malignancy alters the fatty acids of membranes, and fatty acid inputs are known to influence tumorigenesis (15–23).

While MYCN has been implicated in fatty acid uptake, little is known about MYCN or MYC control downstream of *de novo* fatty acid synthesis, which usually terminates with monoenoic oleic acid (OA) when sufficient polyunsaturated FA are available. Unsaturated fatty acid synthesis (UFAS), which is mediated by a subset of FAS genes (**Figure 1b**), has been little studied with respect to MYC and MYCN. UFAS genes modify nascent and dietary fatty acids by both chain elongation and incorporation of double bonds at specific positions within the carbon chain (desaturation) (24). The double bond location from the terminal methyl group along with methylene interruption between double bonds defines fatty acids as omega-3 (ω3), ω6, or ω9 (**Figure 1b, Supplemental Figure S1, Supplemental Table S1**) (25). Highly unsaturated FAs (HUFAs), such as ω6 arachidonic acid (ARA, 20:4), serve as substrates for numerous bioactive oxylipins, many of which are immune-stimulating and pro-inflammatory (13).

The ω3 HUFA docosahexaenoic acid (DHA, 22:6) is critical for normal neuronal health and induces apoptosis in multiple cancer cell types (26–28). Inhibition of FA synthesis reduces NB cell proliferation and xenograft growth (12). Among the seven ELOVL elongase family members collectively responsible for fatty acid chain elongation, ELOVL4 initiates the synthesis of very long-chain HUFA (≥C26) and is highly expressed in the retina and in differentiating neurons (29). MYCN represses *ELOVL4* in poorly differentiated NB, making *ELOVL4* a marker of good prognosis and supporting the idea that FAS and metabolism may be important in NB pathogenesis (30).

Gene expression is highly regulated at both transcriptional and post-transcriptional levels. MicroRNAs (miRNAs) are small, ∼22 nucleotide long single stranded RNAs that regulate mRNAs post transcriptionally through sequence-specific binding to the 3’ untranslated regions (3’UTRs) of target mRNAs. miRNAs have been implicated in cancer as either oncogenic (oncomiRs) or tumor-suppressive (TSmiRs) miRNAs (31). Indeed, altered miRNA function is key to NB development, including a central role for the impairment of the *let-7* miRNA family (32–35). Additional NB-associated TSmiRs have been identified, including *miR-1*, *miR*-*22*, *miR-24*, *miR-34*, *miR-101*, *miR-124*, *miR-140*, *miR-150*, and *miR-204* (33,36–44). miRNAs play roles in most cellular activities, including multiple metabolic processes such as the regulation of glucose metabolism, obesity, and fatty acid oxidation by *let*-7, *mir*-22, and *miR*-33, respectively (45–47). *Let*-7 targets genes throughout the insulin-PI3K-mTOR pathway, including *IGF1R*, *INSR*, *IRS2* and *IGF2BP1*. Overexpression in mice also results in impaired glucose tolerance, demonstrating that miRNAs can regulate entire metabolic pathways *in vivo* by targeting multiple pathway genes (48). While miRNAs have been implicated in a variety of metabolic processes throughout glycolysis, their roles in UFAS pathways are largely unknown (**Figure 1a**) (49,50).

In this study, we aimed to understand the expression patterns and outcome associations of saturated and unsaturated fatty acid synthesis, in poor prognosis NB, as well as fatty acid levels *in vitro* in NB cell lines derived from high-risk tumors. Using complementary analysis of publicly available genomic datasets and cultured NB cell measurements, we examined the potential regulation of UFAS genes through MYCN and MYC activity and posttranscriptional regulation by TSmiRs. Finally, we profiled the levels of ω3, ω6, and ω9 unsaturated FAs in human NB cells. A consistent pattern emerged between high-risk NB outcomes, UFAS gene expression and regulation, and ω9 vs. ω3 and ω6 HUFA lipid profiles. This model provides an organizing principle for future studies to fully understand how the regulation of fatty acid synthesis and metabolism influences NB clinical outcomes, with additional implications for other MYC family driven cancers.

## RESULTS

MYC and MYCN are both known to transcriptionally upregulate multiple genes, contributing to the shift toward anaerobic glycolysis in high-risk cancer, and multiple TSmiRs have been shown to post-transcriptionally down regulate relevant genes (**Figure 1a**). One result of this shift is the shunt of pyruvate towards lactate and citrate production, the latter of which contributes to elevated acetyl-CoA production, which is the fundamental building block of *de novo* FAS (Figure 1b). To investigate the potential involvement of FAS and UFAS (U/FAS) genes in NB, we performed KEGG pathway analysis of the metabolic pathways in high-risk and low-risk NB patients. We observed significant enrichment of the UFAS pathway in high-risk disease. The *de novo* FAS pathway was also elevated, although not significantly (**Figure 2a**). Heatmap analysis of the genes in these two KEGG pathways showed an enriched gene cluster of UFAS genes *SCD*, *FADS2*, and *FADS1,* as well as the *de novo* FAS genes *FASN and ELOVL6,* in high-risk patients (**Figure 2b**). These genes comprise most of the FAS genes outlined in parallel fatty acid metabolism pathways (**Figure 1b**), suggesting that these genes may constitute an important genetic circuit in high-risk disease.

**Figure 2.**
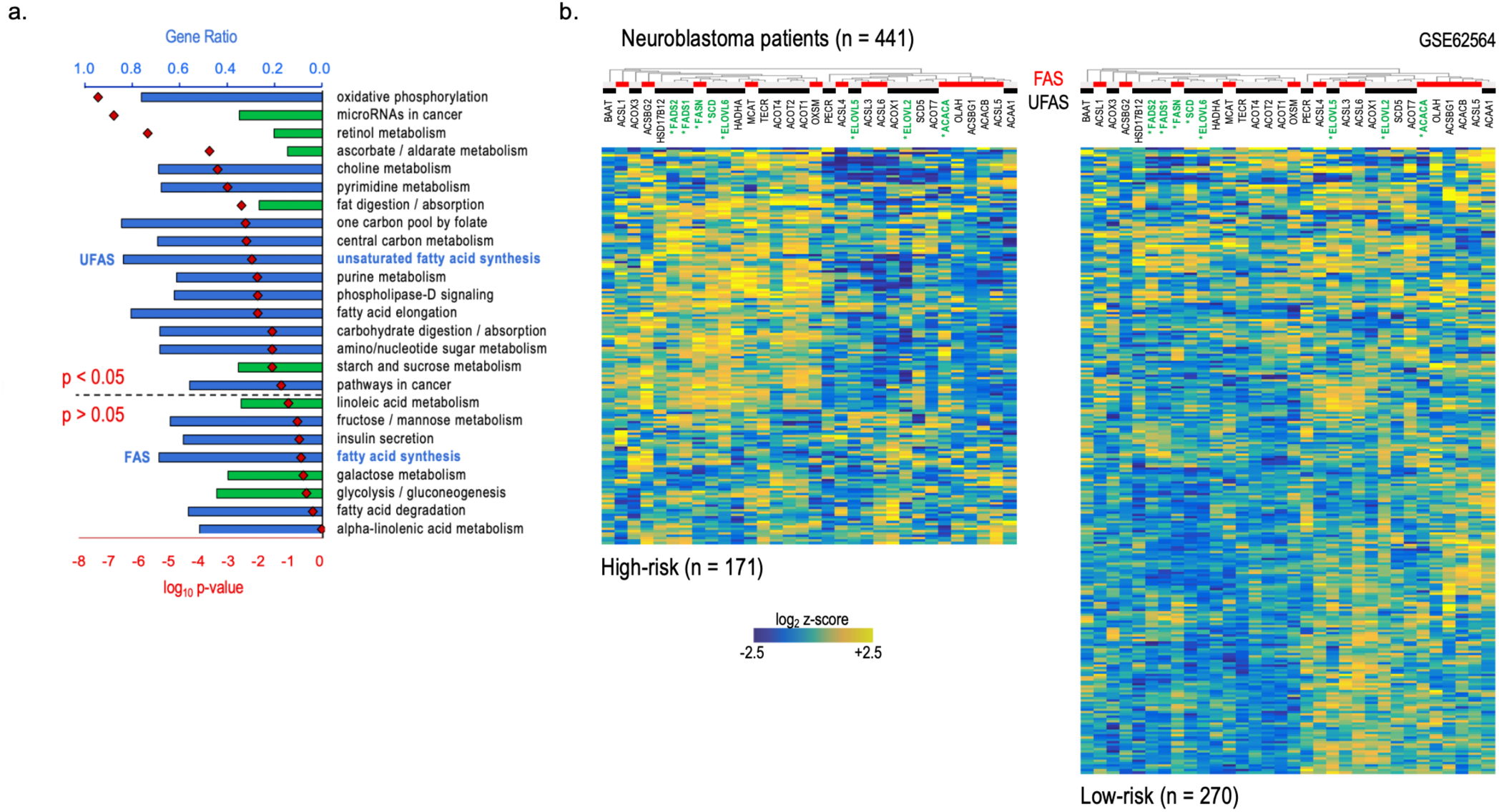
Comparative metabolic KEGG pathway analysis of high and low risk neuroblastoma patients. **a.** KEGG pathway analysis of altered metabolic or cancer related pathways in high-risk (n = 171) and low-risk (n = 270) disease NB patients, stages 1-4. Blue bars are overrepresented pathways; green bars are underrepresented pathways; red diamonds are log_10_ p-values. Over- and underrepresented KEGG pathways were determined by Welch’s t-test followed by B-H FDR correction. (*GEO accession GSE62564*) **b.** Heatmap of unsaturated fatty acid synthesis (UFAS) and fatty acid synthesis (FAS) pathway gene expression in stage 1-4 neuroblastoma patients comparing high-risk (*n=171*) and low-risk disease (*n=270*). U/FAS pathway genes from Figure 1b are titled in green and contain an asterisk (*). Genes associated with FAS and UFAS gene sets are identified by clustered red and black bars, respectively. Expression data presented as normalized values (log_2_ z-scores).

We next compared the expression of U/FAS pathway genes (**Figure 1b**) *ACACA*, *FASN*, *ELOVL6*, *SCD*, *FADS2*, *ELOVL5*, *FADS1*, and *ELOVL2* in high-risk vs. low-risk NB patients. All of these genes, were significantly upregulated in high-risk patients, except *ACACA*, *ELOVL5*, and *ELOVL2* (**Figure 3a**). The upregulated genes were also strongly associated with poor prognosis in NB patients, whereas *ELOVL5* association was not significant. Both *ACACA* and *ELOVL2* were associated with improved prognosis (**Figure 3b**). We also examined the expression patterns of this gene set in eight human NB cell lines derived from high-risk patient tumors using RNA sequencing (RNA-Seq) analysis (**Supplemental Table S2).** We observed that these genes were similarly upregulated compared with normal human fibroblasts (NHF) (**Figure 3c**). *ACACA* was elevated in NB cells vs. NHF, suggesting that *ACACA* may be generally unregulated in NB cell lines compared to normal tissues, but not further enriched in high-risk disease. *ELOVL5* was largely unchanged and *ELOVL2* expression was low overall, which is consistent with human patient expression and overall survival data (**Figures 3a-c**). The expression patterns and prognostic association of these genes in both patients and cell lines suggests that U/FAS synthesis and metabolism play an important role in the pathology of high-risk NB. These observations further suggest that FAS pathway genes *FASN*, *ELOVL6*, *SCD*, *FADS2*, *FADS1* represent a “core” prognostic gene set (U/FAS-core genes) for *de novo* synthesis of ω9 fatty acids and metabolic processing of ω6 and ω3 unsaturated FAs that are uniformly elevated in NB and enriched in high-risk, poor prognosis disease.

**Figure 3.**
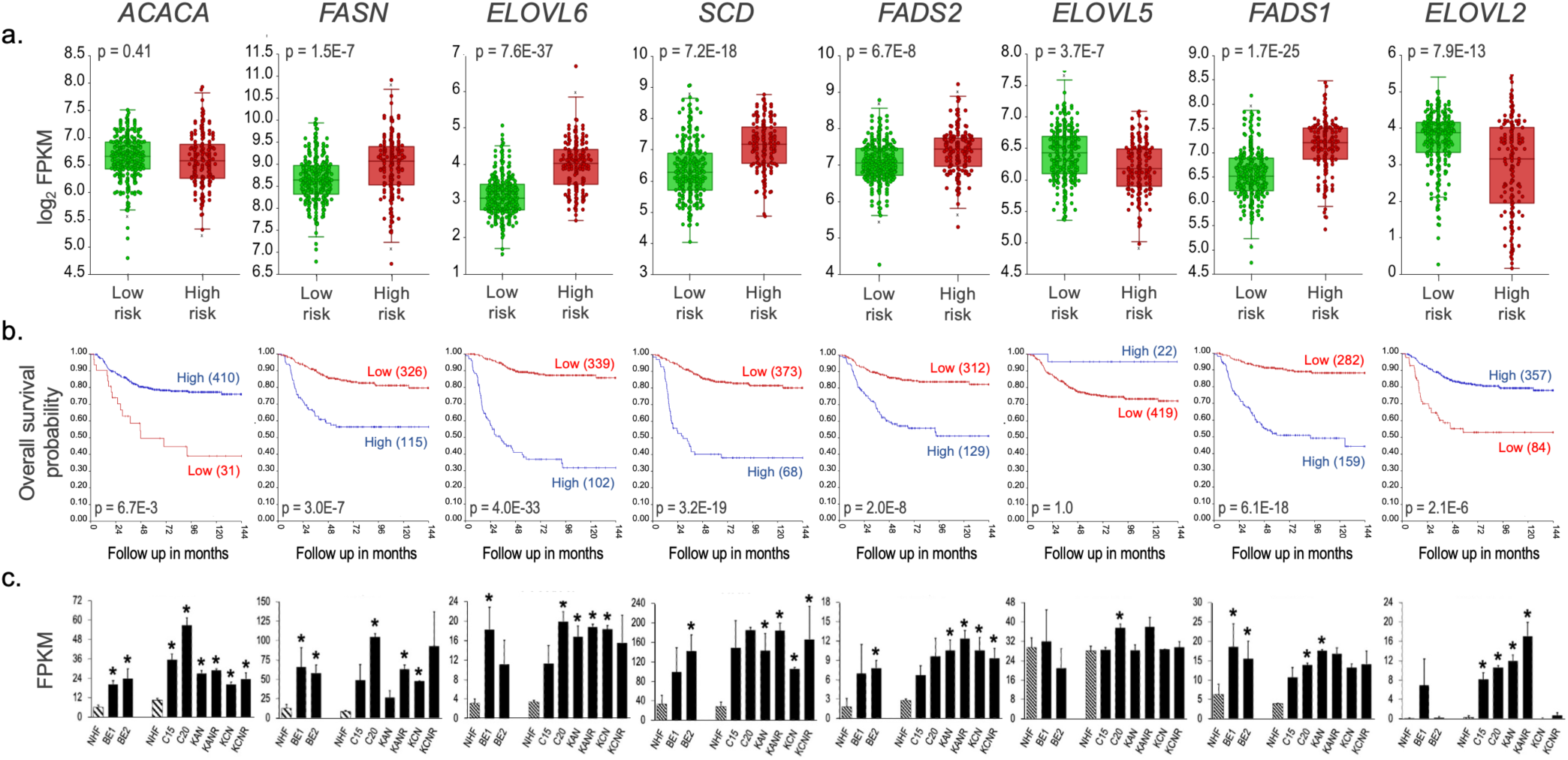
U/FAS genes are elevated in high-risk neuroblastoma and associate with poor prognosis. **a.** Gene expression comparison between NB patients with low-risk (n = 270) and high-risk (n = 171) disease by log_2_ FPKM values. Differential expression significance was determined by one-way ANOVA. **b.** Kaplan–Meier analysis of overall survival in NB patients by gene expression. For each gene, cut-off for high and low gene expression was determined by Kaplan scanning using r2. Significance was calculated using the log-rank test with Bonferroni correction. Number of patients with high and low expression is shown in blue and red parentheses, respectively. **c.** FPKM gene expression levels of U/FAS genes in normal human fibroblasts (NHF) and human neuroblastoma cell lines (left set of NHF, BE1, and BE2 NB cell lines represents n = 3 sequencing rounds; right set of NHF and NB cell lines represents n = 2 sequencing rounds). Significance determined by t-test, \**p*<0.05. (*3a and 3b: GEO accession GSE62564*)

Glycolysis is a highly regulated metabolic process that is controlled by energy requirements, glucose availability, cell state, and hormonal signaling (50). Conversion from oxidative phosphorylation towards lactate production is common in cancer (51). Posttranscriptional regulation of glycolysis pathway genes by multiple TSmiRs can contribute to this conversion (49). In normal tissues and cells, relatively high levels of TSmiRs contribute to glycolysis modulation. In cancer, many of the TSmiRs that negatively regulate glycolysis genes are themselves impaired or downregulated, contributing to the release of most glycolysis genes from normal posttranscriptional inhibition and significantly contributing to a cancer-associated shift away from oxidative phosphorylation-based ATP production (49,52). Therefore, we reasoned that neuroblastoma TSmiRs might similarly target U/FAS pathway genes (53,54). We then analyzed the 3’UTRs of the U/FAS-core gene set for the presence of predicted binding sites for this NB TSmiR list and discovered widespread target-site presence among all core set genes, with an average of 10.4 TSmiR sites per 3’UTR (**Table 1, Supplemental Table S3**). Even the 3’UTRs of U/FAS genes not associated with poor prognosis (*ACACA*, *ELOVL5*, and *ELOVL2*) were highly enriched with TSmiR sites (average of 11.7 sites per 3’UTR), suggesting a unified post-transcriptional regulation of U/FAS pathways by TSmiRs in normal biology and in cancer. MYCN is regulated by TSmiRs in NB and serves as a positive control along with other family members. The *MYCN* 3’UTR also contains nine TSmiR sites, despite its relatively small size. In contrast, the housekeeping genes *SDHA*, *ACTB*, and *GAPDH* have minimal TSmiR sites, with an average of only one site per 3’UTR. Three non-FAS related genes, *DLK1*, *ITGAV*, and *PTEN*, each with long 3’ UTRs, have only 3.7 identified TSmiR sites per mRNA on average. Together, these ratios of TSmiR sites suggest broad post-transcriptional regulation of the U/FAS pathways by TSmiRs, similar to what has been reported for glycolysis. Furthermore, several NB-related TSmiRs, including *miR-1*, *let-7*, *miR-34*, *miR-124*, and *miR-204*, have also been linked to glycolysis gene regulation (**Figure 1a**) (33,36,40,41,44,49,52,55), suggesting that a broad TSmiR regulatory network may influence both glycolysis and U/FAS, and is consistent with a model in which glycolysis and fatty acid synthesis may be co-elevated in cancer, including high-risk NB.

**Table 1.**
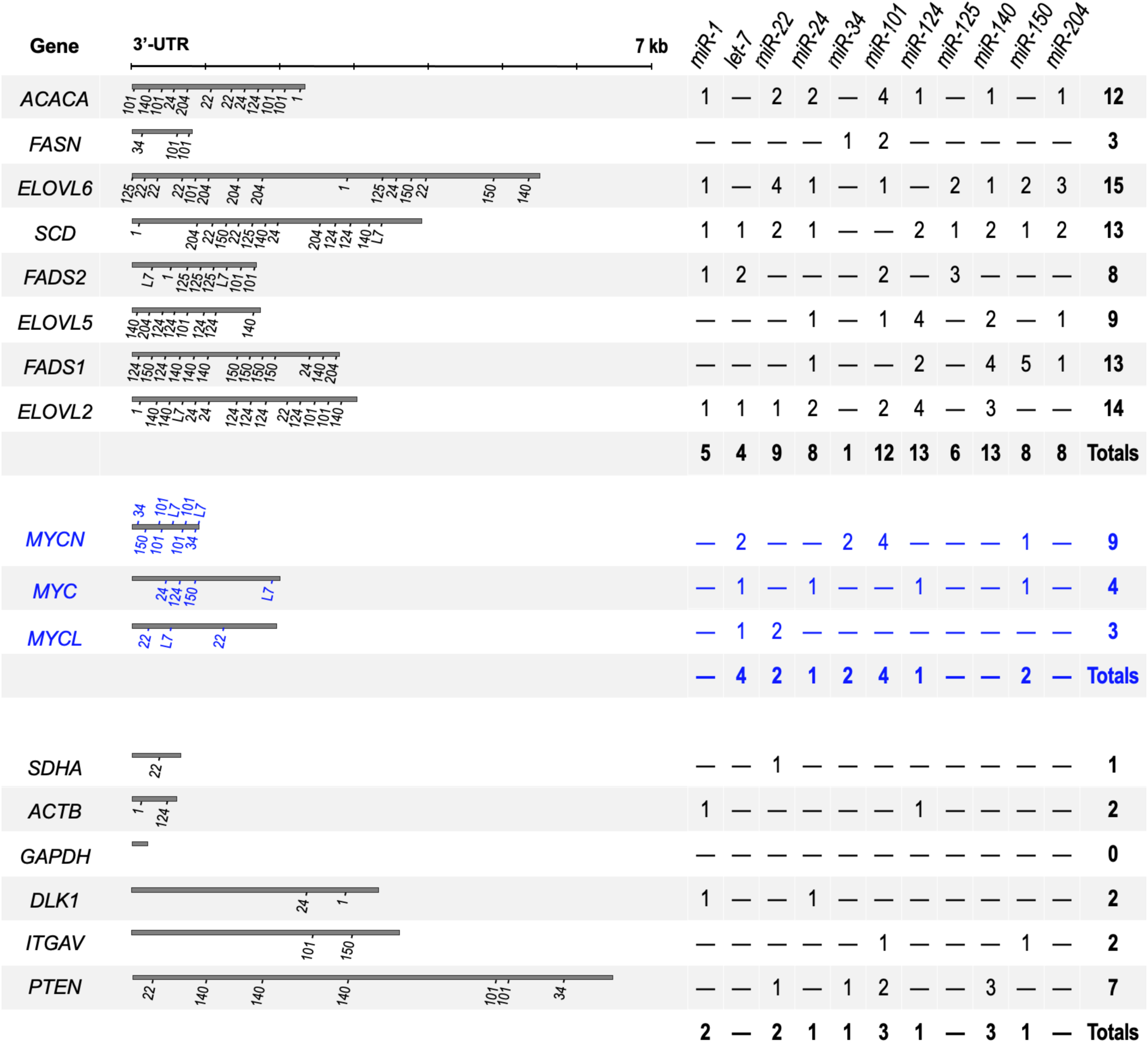
U/FAS gene 3’UTRs showing positions of predicted microRNA target sites. Graphic (left) shows the approximate locations of miRNA target sites within 3’UTRs of listed genes. Scale for 3’UTR length located above UTRs. UTR lengths based on RefSeq sequences for transcript variant 1 in all cases. Numeric table (right) shows the number of sites in multiple U/FAS genes, the *MYC* family, several housekeeping genes (*SDHA*, *ACTB,* and *GAPDH*), and three unrelated genes (*DLK1*, *ITGAV*, and *PTEN*). (*miRNA sites identified through Targetscan, v8.0)*

The TSmiRs assessed here have been previously demonstrated to possess anti-growth, tumor suppressor functions and frequently reduced expression in NB (53,54). We observed their collective downregulation in tyrosine hydrolase-MYCN (*TH-MYCN)*-driven murine NB compared to the wild-type sympathetic ganglia tissue of origin, demonstrating broad inhibition of multiple TSmiRs during NB development and collectively confirming previous reports of impaired TSmiR expression and function in NB (**Figure 4a**) (53,56,57). The oncogenic *miR-17-92* cluster (which includes miRNAs *miR-17*, *miR-18a*, *miR-19a, miR-19b*, *miR-20a*, and *miR-92a*) is a known transcriptional target of MYCN and is upregulated in *TH*-*MYCN* tumors, as expected.

**Figure 4.**
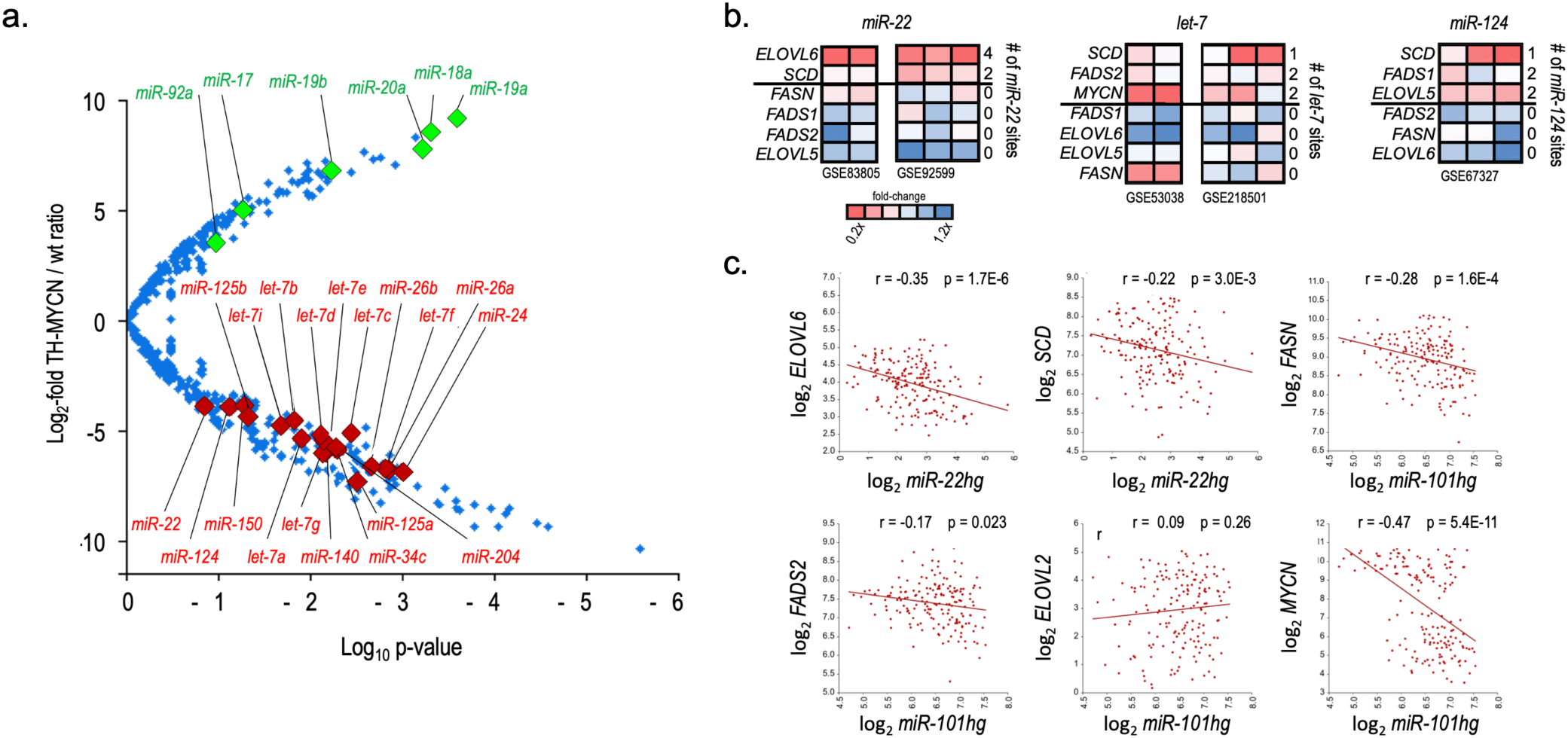
Tumor suppressor miRNAs are downregulated in TH-MYCN NB and result in reduced expression of their U/FAS mRNA targets. **a.** Volcano plot of differentially expressed miRNAs in *TH-MYCN* NB tumors (n = 12) and wildtype sympathetic ganglia (n = 12; NB tissue-of-origin). The x-axis indicates log_10_ p-values; y-axis indicates log_2_-fold change between tumors and ganglia. Red diamonds: tumor suppressor miRNAs from *Table 1*; green diamonds: miRNAs of the miR-17-92 oncomiR cluster (*r2 dataset identifier E-MTAB-2618*). **b.** Heatmaps of U/FAS genes expression changes in response to *miR-22* (left), *let-7* (middle), or *miR-124* (right) overexpression. Each heat map is a unique study labeled with its GEO accession number. Each column represents a replicate experiment. Number of miRNA sites identified in the 3’UTR of the listed gene are listed along the right of each miRNA set. **c.** Correlation plots for mRNA-target gene expression *vs.* miRNA-host gene for *miR-22* or *miR-101* in high-risk NB patients (n=176). (*5c: GEO accession GSE62564*)

A recent meta-analysis of miRNA-mRNA inhibition mechanisms revealed that at least half of all miRNA target genes are primarily regulated through mRNA decay (58). We thus reasoned that mRNA levels of U/FAS TSmiR target genes may be reduced by elevated TSmiR activity; therefore, we examined the effects of TSmiR overexpression on *miR-22*, *let-7*, and *miR-124* targeted FAS mRNA levels in existing TSmiR overexpression datasets (**Figure 4b**). Fold-change heatmaps of two independent *miR-22* overexpression studies displayed similar patterns of reduced expression of *miR-22* targets *SCD* and *ELOVL6,* which contains four miR-22 sites in its 3’UTR (**Table 1**) and was strongly suppressed in both studies. In contrast, the expression levels of both *FADS2* and *ELOVL5*, which were not targeted by *miR-22*, were not affected (**Figure 4b, left panel**). A similar pattern of reduced expression was observed for *let-7* targets *SCD*, *FADS2*, and *MYCN* in studies of de-repressed *let-7* biogenesis and *let-7* transfection. Levels of non-*let-7* targets *FADS1*, *ELOVL6*, and *ELOVL5* were relatively unaffected, whereas non-*let-7* target *FASN* levels were reduced in the *let-7* biogenesis study but minimally affected by transfected *let-7* (**Table 1**; **Figure 4b, middle panel**). *MiR-124* transfection resulted in the reduction of miR-124 targets *SCD*, *FADS1*, and *ELOVL5*, whereas none of the non-miR-124 targets *FADS2*, *FASN*, or *ELOVL6* were decreased (**Table 1**; **Figure 4b, right panel**).

miRNA genetic loci are often located within the introns of larger host genes (hg) and are frequently coordinately transcribed as part of the host transcript from which they are processed (59,60). Thus, miRNA host gene expression patterns can serve as a surrogate for intragenic miRNA expression levels when small RNA sequencing data or qPCR analyses are not available. Therefore, we compared the expression levels of *miR-22hg* and *miR-101hg* to the expression levels of the individual *miR-22* targets *ELOVL6* and *SCD*, and *miR-101* targets *FASN*, *FADS2*, *ELOVL2*, and *MYCN* (**Figure 4c**). Both *miR-22* target genes were inversely correlated with the expression of the miRNA host genes. *FASN*, *FADS2,* and *MYCN* were inversely correlated with *miR-101hg* expression, whereas *ELOVL2* expression was not significantly correlated. These expression patterns provide further support for a model in which U/FAS genes are broadly targeted by TSmiR in NB. While these results are consistent with the targeting of FAS genes by TSmiRs, further work is required to validate these observations.

Overexpression of both *MYCN* and *MYC* drives metabolic conversion to favor the rapid generation of ATP via anaerobic glycolysis over oxidative phosphorylation (9,10,30). While *MYCN* is the dominant oncogene in NB, MYC has been reported to play a role in some *MYCN* non-amplified tumors (6). Given the strong overlap in MYCN and MYC functions, we next examined the relative expression of *MYCN* and *MYC* in both *MYCN*-amplified and non-amplified disease in three distinct NB patient studies. *MYCN* exhibited the expected high expression pattern in amplified disease across all three patient datasets, displaying remarkable consistency in expression levels in both MYCN-amplified and non-amplified tumors (**Figures 5a, 5b**). *MYC* also displayed stable but lower expression patterns across the datasets, revealing a reciprocal expression pattern with *MYCN* (**Figures 5a, 5b)**. When considered together, total *MYC/N* expression levels were higher in all patients from all three studies than in tissues of origin adrenal gland and trunk neural crest, demonstrating that even non-amplified tumors have an elevated *MYC/N* signature. *MYCN* amplified patients had the highest *MYC/N* dose, as expected, although the magnitude of *MYCN* increase over non-amplified is likely somewhat mitigated by contributing yet reciprocal *MYC* expression. This cooperative pattern is consistent with MYCN and MYC having similar effects on metabolic reprogramming.

**Figure 5.**
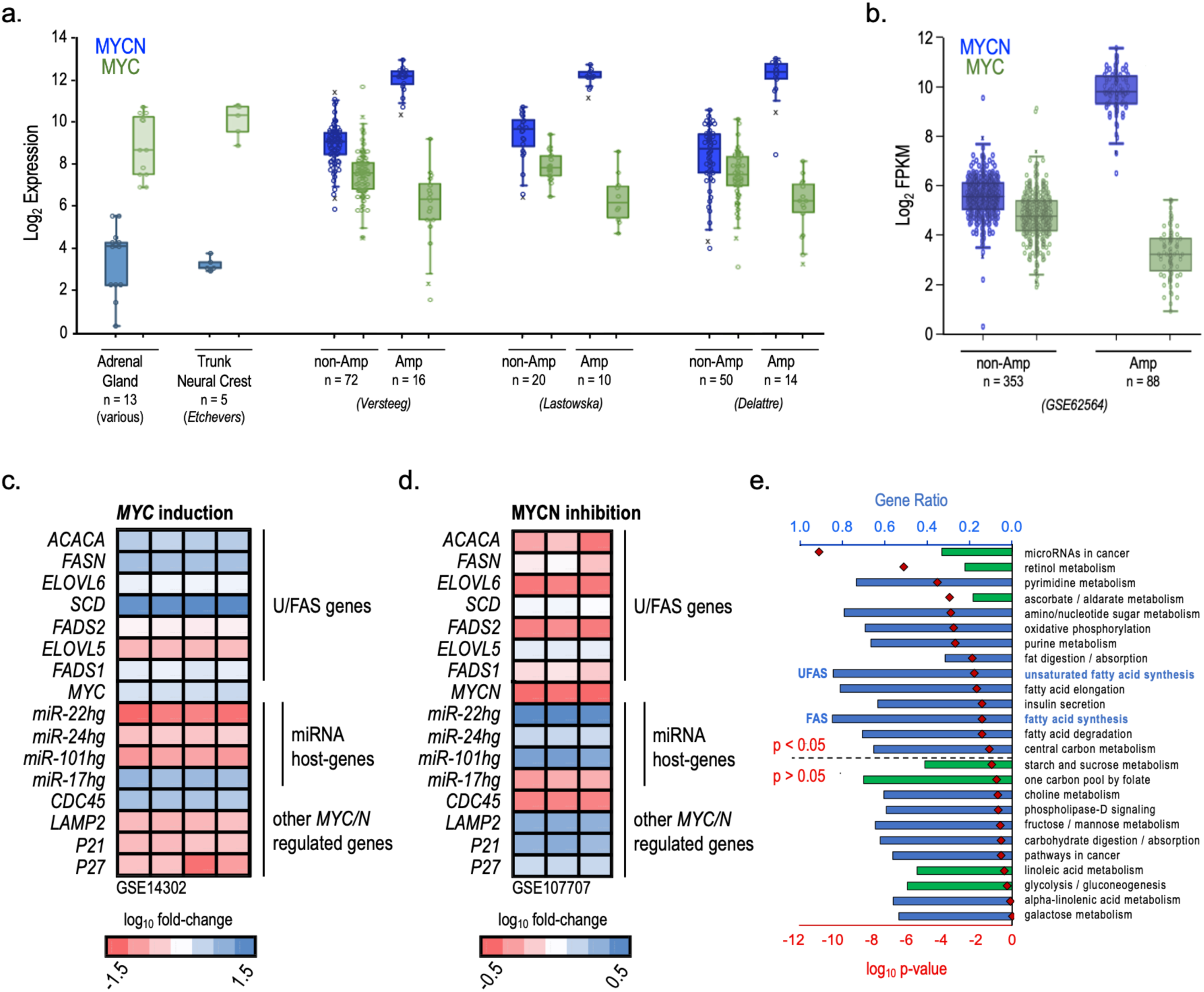
Expression patterns of MYC/N and related genes in NB by KEGG Gene Set Analysis. **a.** Microarray expression of *MYC* and *MYCN* in MYCN-amplified and non-Amplified patients from three different NB data sets and normal Adrenal Gland (NB tissue of origin) and Trunk Neural Crest cells (NB cells of origin). **b.** Gene expression comparisons in log_2_ FPKM of *MYC* and *MYCN* in MYCN-amplified (n = 88) and non-amplified (n = 353) NB patients. **c-d.** Heatmaps of U/FAS and MYC/N target gene expression after MYC induction *(c)* or MYCN inhibition by JQ-1 bromo-domain inhibitor *(d)*. Each column represents a replicate experiment. **e.** KEGG pathway analysis of altered metabolic or cancer related pathways in *MYCN*-amplified (n = 88) vs. non-amplified (n = 353) NB patients, stages 1-4. Blue bars are overrepresented pathways; green bars are underrepresented pathways; red diamonds are log_10_ p-values. Over- and underrepresented KEGG pathways were determined by Welch’s t-test followed by B-H FDR correction. (*5e: GEO accession GSE62564)*

Given the predominance of *MYC/N* levels in amplified NB and the strong connection between the *MYC* family and metabolic reprogramming, we hypothesized that both *MYC* and *MYCN* would also influence U/FAS gene expression. Therefore, we analyzed U/FAS gene expression in complementary datasets of induced *MYC* expression in P493-6 lymphoid cells (**Figure 5c**) and JQ-1 bromodomain inhibitor-mediated MYCN inhibition in BE(2)C NB cells (**Figure 5d**) (61). As previously shown, *MYC* induction resulted in a significant upregulation of the U/FAS genes *ACACA* (6.6 x), *FASN* (9.2 x), and *SCD* (33.2 x), and *ELOVL6* (2.6 x). *FADS2*, *ELOVL5*, and *FADS1* expression levels were also significantly increased by 2.6 x, 1.6 x, and 3.1 x, respectively. Known MYC-activated (*MIR17HG* and *CDC45*) and suppressed (*LAMP2*, *P21*, and *P27*) genes behaved as expected. The host genes for *miRNA-22*, *miRNA-24*, and *miRNA-101* were also suppressed, suggesting that MYC may directly inhibit their transcription (**Figure 5c**). In BE(2)C cells treated with JQ-1, U/FAS genes were consequently downregulated between 0.4 x and 0.67 x (**Figure 5d**). *FADS2* and *ELOVL6* were the most affected genes, with expression decreasing to 33% of the control in both cases. MYC family target genes once again behaved as expected, with decreased *MIR17HG* and *CDC45* levels and de-repressed *LAMP2* and *P21* expression. Consistent with *MYC* induction, inhibition of MYCN resulted in increased levels of host genes for *miRNA-22*, *miRNA-24*, and *miRNA-101*, further supporting a repressive role of MYC/N in their regulation.

These data suggest that both MYCN and MYC broadly regulate the U/FAS pathway, identifying novel roles for MYC/N transactivation of *FADS2* and *FADS1*, with *ELOVL6* as an additional novel target of MYCN. Furthermore, KEGG analysis of *MYCN*-amplified vs. non-amplified NB patients showed that several fatty acid pathways were significantly enriched in the *MYCN*-amplified group (**Figure 5e**). Half of high-risk NB patients are *MYCN*-amplified and had a significantly enriched FAS pathway. Among *MYCN*-amplified patients, KEGG pathways for FAS, fatty acid elongation, fat digestion/absorption, and fatty acid degradation were all significantly elevated, further supporting MYCN regulation of U/FAS metabolism in NB. miRNA host genes *miR-22hg, miR-24hg, and miR-101hg* appeared to be suppressed by both MYCN and MYC modulation (**Figures 5c, 5d**), providing a possible mechanism for TSmiR suppression in NB and perhaps other MYC-driven cancers. It is also possible that U/FAS gene expression levels in cancer are, in part, regulated indirectly by MYC and MYCN through transcriptional suppression of TSmiR host gene expression and consequent release of U/FAS mRNAs from posttranscriptional TSmiR-mediated degradation. This model establishes a novel regulatory network between *MYC*/*N*, U/FAS core genes, and multiple TSmiRs (which also target *MYCN* mRNA in NB).

To determine whether the observed expression levels of FAS genes correlates with changes in cellular fatty acid profiles, we analyzed eight NB cell lines using gas chromatography and mass spectrometry (**Figure 6a, Supplemental Tables S1 and S2**). Levels of ω9 Mead acid and its elongation product, docosatrienoic acid (DTrA), were dramatically increased beyond levels seen in normal cells (**Figure 6a, Supplemental Table S4**). Although grown under the same conditions, NHF had a higher proportion of ω3 DHA than all NB cells. Elevated Mead acid (20:3ω9), normalized to arachidonic acid (ARA, 20:4ω6), known as the triene: tetraene (T:T) ratio, is the accepted biochemical index of essential fatty acid deficiency (EFAD) (62). In plasma, a T:T greater than 0.4 indicates EFAD; a similar threshold applies to cells (63). In NB cell lines, ARA levels decreased from 10% w/w in NHF to ∼5% w/w, while Mead acid increased from trace levels to 2% - 5% of total fatty acids (**Figure 6a, Supplemental Figures S2a and S2b**), reflecting EFAD for all cells. The sum of Mead acid and ARA was 8.8%±1.2%, w/w (mean ± SD), reflecting the compensatory increase in Mead acid levels as ARA decreased (**Supplemental Figures S2c and S2d**) (63). Normally, EFAD arises because of the restricted supply of the ω6 essential FA linoleic acid (**LA**) (**Figure 1b, Supplemental Table S1**) (64). However, NHF grown in the same media as the NB cells had trace amounts of Mead acid and thus were not EFAD. The enhanced expression of U/FAS genes compared to NHF cells (**Figure 3c**), as well as the pattern of fatty acid levels discussed below, points to unusually high synthesis of ω9 HUFA, and possibly Mead acid specifically.

**Figure 6.**
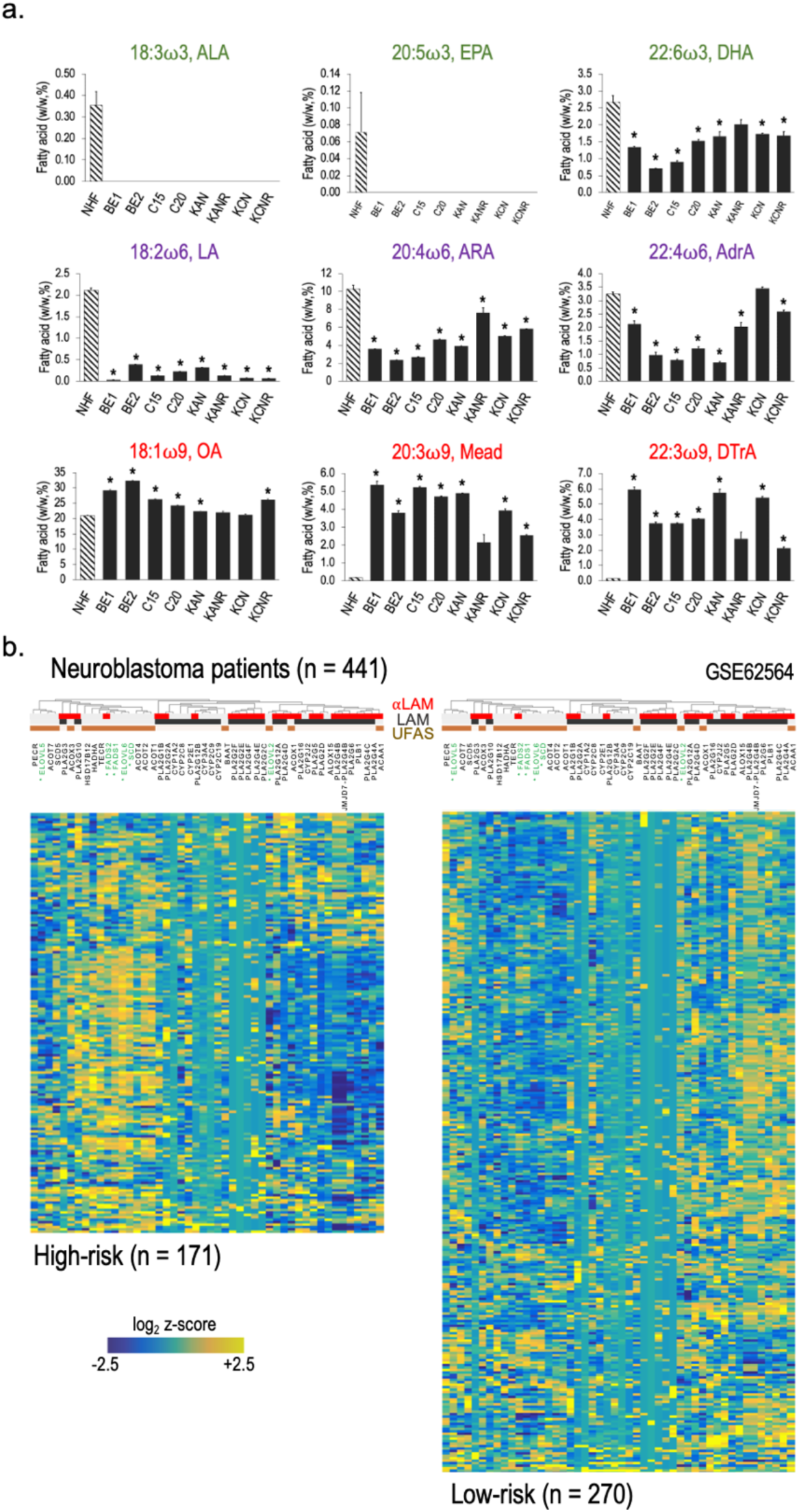
Omega-9 unsaturated fatty acids are elevated in NB. **a.** Fatty acid levels in normal human fibroblasts (NHF; striped bars) and human NB cell lines, presented as percent of overall lipid weight (w/w, %). Pathway precursors and key HUFAs are shown. Omega-3 fatty acids are titled in green, omega-6 are titled in purple, and omega-9 fatty acids are titled in red. **b.** Heatmap of ⍺-linolenic acid metabolism (αLAM), linoleic acid metabolism (LAM), and unsaturated fatty acid synthesis (UFAS) gene expression comparing high-risk (n = 171) and low-risk (n = 270) disease in NB patients, stages 1-4. U/FAS pathway genes from figure 1b are titled in green and contain asterisks (*). Genes associated with αLAM, LAM and U/FAS gene sets are identified by clustered red, black, and brown bars, respectively. Expression data is presented as normalized values (log_2_ z-scores).

KEGG pathway analysis of FAS pathways in high-risk and low-risk NB patients revealed that while the *de novo* FAS pathway genes were uniquely enriched in high-risk patients, the majority of ω3 precursor ALA and ω6 precursor LA metabolism genes, represented by UFAS were suppressed. In contrast, genes from the three pathways were largely uniformly expressed in both the normal adrenal gland and tibial nerve tissue samples (**Figure 6b, Supplemental Figure S3**), suggesting that a genetic program favoring *de novo* FAS over UFAS may be selected for high-risk NB.

HUFAs are precursors of eicosanoids and other locally active bioactive signaling lipids. Derivatives of ω6 ARA are produced via three main pathways driven by lipoxygenases (e.g., ALOX5), cyclooxygenases (e.g., COX2), and CYP450 enzymes (**Figure 7a**) (13). Of the more than 50 unique CYP450 enzymes, 15 are epoxygenases that act on HUFAs such as ARA. Only two of these 15 are expressed in NB; *CYP2C8* and *CYP4V2* (**Supplemental Figure S4a**). ARA processing by these enzymatic arms produces several pro-inflammatory eicosanoids, including prostaglandins, thromboxanes, leukotrienes, EETs, and 20-HETE. These molecules can stimulate immune activity and are involved in tumor formation and progression (65). *ALOX5* is the most abundant of the six human lipoxygenases in NB and has been implicated as a tumor suppressor (**Supplemental Table S4b**) (66). The enzymes producing these pro-inflammatory eicosanoids are uniformly down regulated in high-risk NB, and low expression of this gene set is strongly associated with poor prognosis (**Figures 7b, 7c**), suggesting a selective advantage for reduced ω6 levels in NB and is consistent with what we observed in NB cell lines.

**Figure 7.**
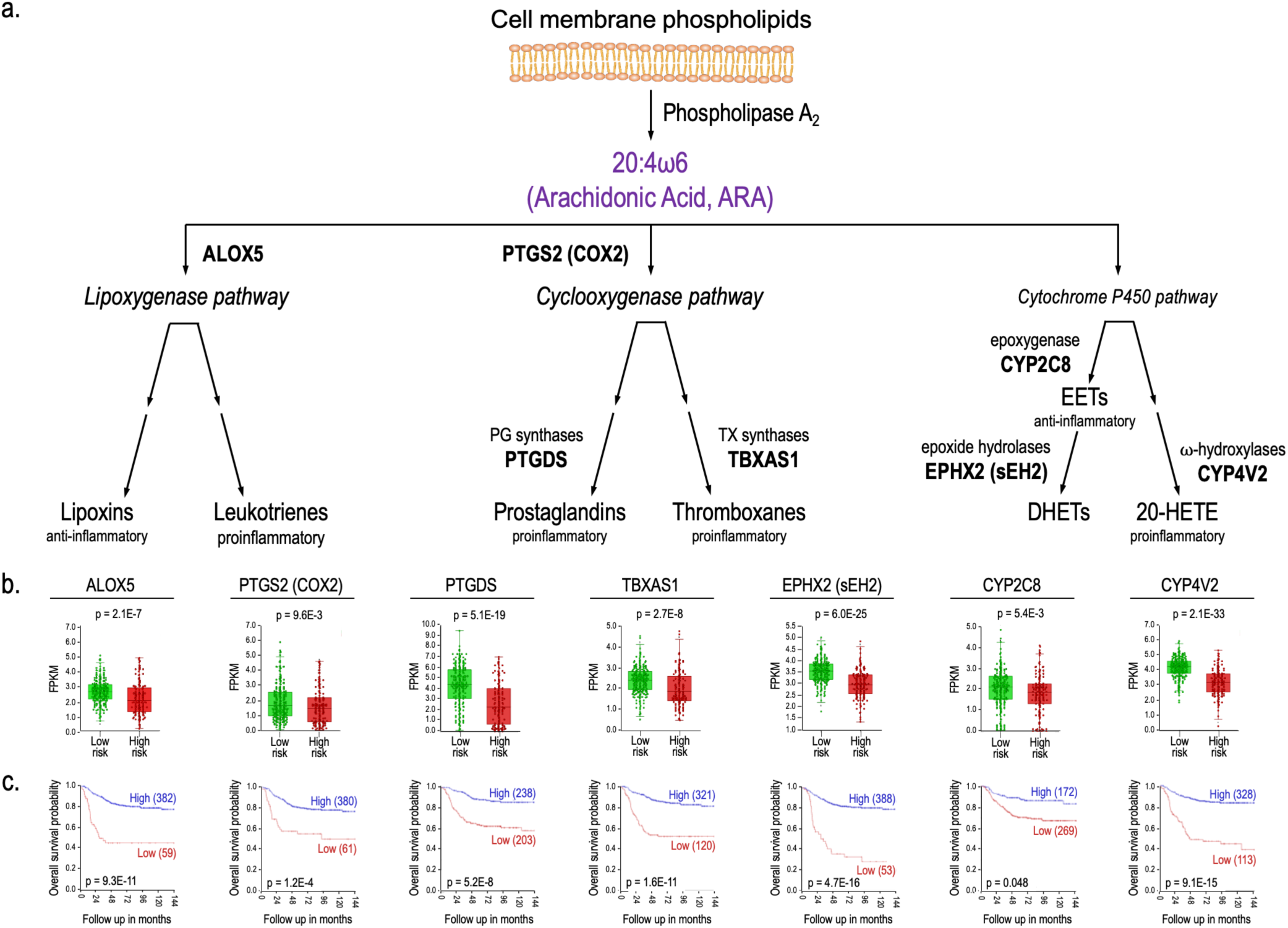
ARA pro-inflammation metabolism pathways are downregulated in high-risk neuroblastoma. **a.** Schematic of ARA immune-active metabolism. ARA, 20:4μ6 is metabolized by the lipoxygenase, cyclooxygenase, and cytochrome P450 enzymatic pathways to produce various immune active eicosanoids. **b.** Associated gene expression comparison between NB patients (stages 1-4) with low-risk (n = 270) and high-risk (n = 171) disease by RNA-sequencing FPKM values. Differential expression significance was determined by one-way ANOVA. **c.** Overall NB survival plots of ARA metabolizing enzymes are located next to genes of interest. Number of patients with high and low expression is shown in blue and red parentheses, respectively. Cut-off for high and low gene expression for Kaplan-Meier analysis was determined by Kaplan scanning using r2. Significance was calculated using the log-rank test with Bonferroni correction. (*7b and 7c: GEO accession GSE62564*)

## DISCUSSION

*MYC*, *MYCN*, and numerous TSmiRs have been implicated in glucose metabolism. We have shown here that the MYCN/TSmiR regulatory program is similarly involved in U/FAS pathway regulation in NB with elevated expression of U/FAS-core genes, *FASN*, *ELOVL6*, *SCD*, *FADS2*, and *FADS1* in high-risk disease, which is strongly associated with poor prognosis (**Figures 1-3**). Remarkably high levels of ω9 HUFA Mead acid and DTrA in cultured NB cells and concordant gene expression data between cells and NB patients suggest that the upstream FAS pathway consisting of FASN and ELOVL6 can initiate the U/FAS anabolic pathway for the directed production of ω9 unsaturated FAs. Careful inspection of the ω3 fatty acids showed that only terminal ω3 DHA was present at an appreciable concentration, with precursor ALA and intermediate ω3 EPA at trace levels (**Figure 6a**). Similarly, for ω6 fatty acids, ω6 precursor LA is low in all cells, while the usual terminal product ω6 ARA and its elongation product ω6 AdrA are present. Indeed, there is some indication that ω9 OA may be elevated beyond its already dominant level in some NB cell types. Upregulation of the U/FAS genes and the upstream pathway may induce superficial essential fatty acid deficiency via unusually high production of Mead acid.

We propose a model in which MYCN and TSmiRs oppositely regulate fatty acid synthesis in NB as an extension of the MYC-driven Warburg effect, where shifting ATP production from oxidative phosphorylation towards glycolysis connects directly to unsaturated fatty acid synthesis through the increased production of acetyl-CoA, the fundamental building block of *de novo* FAS. This model introduces the UFAS genes *ELOLV6*, *FADS2*, and *FADS1* as novel targets of MYCN in NB and the broad collective posttranscriptional regulation of FAS and UFAS genes by multiple TSmiRs (**Figure 4**). This model has implications for MYC family-driven cancers (**Figure 5**).

While reports of ω9 Mead acid in cancer are generally scarce, it was shown to be elevated in both the serum and tumors of 38% of patients with hepatocellular carcinoma. It was not present in other tissues, suggesting that Mead production was elevated only in the tumor itself (22). Mead acid treatment increased proliferation of HRT-18 breast cancer cells, whereas ω3 EPA (**Figure 1b, Supplemental Table S1**) reduced cell growth (67). Furthermore, Mead acid increases both invasion activity and cell growth of in multiple squamous cell cancer lines (68). Together, these studies suggest that Mead acid may provide a growth advantage in cancer, which could provide a selective rationale for elevated ω9 Mead acid production in NB.

*FADS2* dysregulation at the 11q13 major cancer hotspot region alters fatty acid metabolism in several cancer types (69). FADS2 operates on at least 16 substrates, catalyzing the biosynthesis of several unsaturated fatty acids by Δ6, Δ8, and Δ4 desaturation (24). The availability of fatty acid substrates for FADS2 in any tissue is the major determinant of the final product mixture for further membrane lipid synthesis and/or synthesis of signaling molecules (70–72). *FASN*, *ELOVL6*, and *SCD* comprise the classical *de novo* FAS pathway and produce ω9 oleic acid (OA) (**Figure 1b**). These three genes were also among the most upregulated genes in the U/FAS-core gene group, which is consistent with high OA production (**Figures 3a, 3c**). Indeed, the actual amount of OA produced is likely even higher than observed given that OA is further processed to downstream ω9 fatty acids, such as Mead acid and DTrA. (**Figures 1b, 6a**).

Our observations outline a possible genetic disposition towards lowering ω3 and ω6 fatty acid metabolism in high-risk NB. Genes from the ALA (ω3) and LA (ω6) metabolism pathways were down regulated in high-risk disease compared to the UFAS-core genes *FASN*, *ELOVL6*, *SCD*, *FADS2*, and *FADS1*, which were significantly enriched in high-risk over low-risk disease and in *MYCN* amplified over *MYCN* non-amplified NB (**Figure 2, 6b**). The FADS1 and FADS2 desaturases introduce double bonds into ω3, ω6, and ω9 fatty acid chains. They have the highest affinity for ω3 fatty acids, followed by ω6 and then ω9, and yet ω9 HUFAs were enriched in NB cells (**Figure 6a**). Cellular ω3 ALA and ω6 LA precursors were depleted to trace levels, thus unmasking the *de novo* synthesis of Mead acid and DtrA. The production of Mead acid and DTrA in NB may be driven by significantly increased ω9 substrate availability, as well as upregulated UFAS genes. FASN, ELOVL6, and SCD complete *de novo* FAS and produce oleic acid (OA), which is the first ω9 fatty acid in the ω9 pathway (**Figure 1b**). These three genes were also among the most upregulated genes in the the UFAS-core group, suggesting that flux through OA production may be high (**Figure 3a**).

Selective pressures may also explain the decreased ω3 and ω6, and elevated ω9 and FA levels observed in NB (**Figure 6a**). Multiple studies have shown that ω3 HUFAs EPA and DHA have anticancer activity through the induction of apoptosis in multiple cancers (**Figure 1b; Supplemental Table S1**) (26–28). DHA treatment of ovarian and lung cancer cell lines and an *in vivo* mouse model of ovarian cancer increased reactive oxygen species (ROS) production, induced apoptosis *in vitro,* and reduced Ki67 staining and tumor size *in vivo* (73,74). Interestingly, pre-treatment with the antioxidant NAC reversed these effects, suggesting that ROS production may be a key mechanism of growth inhibition by DHA. DHA and other HUFAs are known to be highly sensitive to ROS, which attack double bonds in fatty acid chains. ROS/HUFA reactions produce hydroperoxides on fatty acid chains, resulting in chain reactions that further propagate ROS production and damage (75). These observations and our own provide a plausible rationale for gaining a selective advantage by suppressing ω3 synthesis in NB.

Chronic inflammation is associated with increased cancer incidence, whereas an inflammatory tumor microenvironment and acute inflammation have been shown to increase anti-cancer immune function and immune therapy efficacy (76). Local inflammation has also been linked to immune cell infiltration. While some adult tumors, such as melanoma, are considered “hot” tumors with high local inflammation and immune cell presence, pediatric solid tumors are most often “cold” (77). NB is a classic cold tumor and was recently reported to have the lowest inflammatory response signature of 37 tumor types, demonstrating that NB strongly suppresses inflammatory signaling. *MYCN* amplification was shown to further reduce inflammatory signaling, which is consistent with the limited efficacy of immune therapies in NB (78).

Our observations regarding ω6 fatty acids provide a plausible explanation for the reduced inflammatory signature in NB. Both LA and ARA serve as key precursors of pro-inflammatory eicosanoids (**Figures 1b, 1c**) (13). Lower levels of both LA and ARA in NB reduce the pool of precursors available for production. We further observed reduced expression of all three (*LOX, COX, and Cytochrome P450*) enzyme pathways that produce inflammatory eicosanoids from ARA in high-risk NB, and found that low expression of genes in these pathways was strongly associated with worse prognosis (**Figure 7**). Our observations are consistent with the previously established low inflammation signature in NB and provide a credible model for metabolic reprogramming that favors ω9 fatty acid production over ω6 ARA production. Furthermore, recent studies have also demonstrated the pro-apoptotic effect of ARA on cancer cells and revealed the anti-tumor effects of ARA metabolism, providing further support for selective pressure to minimize both ARA and downstream eicosanoid production(79–82).

Here, we show that fatty acid synthesis is deregulated in high-risk NB by both MYCN activity and TSmiR disruption and contributes to poor outcomes. We also reported altered lipid profiles in NB cells by decreasing ω3 and ω6 and increasing ω9 fatty acid levels, which may be selected for to minimize ω3-driven ROS toxicity and ω6-driven pro-inflammatory signaling. Our results suggest a model of unified metabolic reprogramming of unsaturated fatty acid synthesis that may be a novel extension of the Warburg effect, with a series of implications for MYC family driven cancers, mechanisms for suppression of inflammation in cold tumors, and new opportunities for therapies that modulate dietary intake and synthesis of unsaturated fatty acids.

## MATERIALS AND METHODS

### Cell Lines

The following cell lines were used in this study: NHF (BJ, RRID CVCL_3653), BE1 (SK-N-BE(1), RRID CVCL_9898), BE2 (SK-N-BE(2), RRID CVCL_0528), C15 (CHLA-15, RRID CVCL_6594), C20 (CHLA-20, RRID CVCL_6602), KAN (SMS-KAN, RRID CVCL_7131), KANR (SMS-KANR, RRID CVCL_7132), KCN (SMS-KCN, RRID CVCL_7133), KCNR (SMS-KCNR, RRID CVCL_7134). All NB cell lines were obtained from the Childhood Cancer Repository at Texas Tech University Health Science Center, which is part of the Children’s Oncology Group. BJ normal human fibroblasts (NHF) were purchased from ATCC and used as normal control cells. All these cells were grown in RPMI-1640 medium supplemented with 10% fetal bovine serum and were tested for mycoplasma before and during use. Details for all the cell lines used in this study are shown in *Supplemental Table S2*.

### RNA-Seq

Total RNA was submitted to LC Sciences (Houston, TX) for library prep and sequencing. In the present study, we report UFAS genes and will report other genes elsewhere. RNA-seq data sets are available under the GEO reference series GSE (*to be deposited*) (http://www.ncbi.nlm.nih.gov/geo/query/acc.cgi?acc=GSE(TBD)). Two batches of NB cell lines were subjected for RNA-Seq analysis; Batch 1: NHF, BE1, and BE2 (three biological replicates per cell line), Batch 2: NHF, C15, C20, KAN, KANR, KCN, and KCNR (two biological replicates per cell line).

### microRNA target site identification

The TargetScan platform (version 8.0) was used to identify predicted miRNA target sites within the 3’UTR for each gene using the RefSeq sequence from transcript variant 1 (83). For each gene, TargetScan lists all predicted miRNA sites within the 3’UTR region of a transcript pertaining to a single Ensemble ID (ENST). For some of the genes the Ensembl 3’UTR transcript analyzed by TargetScan was longer than that found within the RefSeq sequences for a gene’s transcript variant 1. We adhered to the RefSeq 3’UTR length (*Table 1*). Briefly, 3’UTRs were matched to the predicted binding of TSmiRs known to be associated with NB, specifically *miR-1*, *let-7*, *miR-22*, *miR-24*, *miR-34*, *miR-124, miR-125*, *miR-150,* and *miR-204*.

### Fatty Acid Profiling

Fatty acid analysis of cultured cells was performed as described previously (72,84–86). Briefly, cell pellets were harvested after centrifugation and fatty acid methyl esters (FAME) were prepared according to a modified one-step method (three biological replicates per cell line) (87). FAME were structurally identified by gas chromatography (GC) - chemical ionization mass spectrometry (GC/CI-MS/MS) (88,89) and quantified using a GC-flame ionization detector (GC-FID) after calibration with an equal-weight FAME standard mixture to develop response factors. The statistical tests used are shown in each figure legend. Statistical significance was set at *p* < 0.05.

### Secondary analysis

The SEQC-498 human NB patient dataset from “An Investigation of Biomarkers Derived from Legacy Microarray Data for Their Utility in the RNA-Seq Era” was used for human NB patient analysis (90). Gene expression graphs, Kaplan-Meier curves, and KEGG pathway analysis of human NB patients were generated using the data analysis tool suite on the R2: Genomics Analysis and Visualization Platform (91). Other datasets analyzed on the R2 database: For analysis of miRNA expression in TH-MYCN murine NB (*Figure 4a*): “The role of miRNAs in NB tumor development” (R2 internal identifier: mir_avgpres_thmycn24_mirbase19mm2). For analysis of MYCN and MYC expression across NB patient datasets (*Figure 5a*): Control tissues; Adrenal Gland (various) (R2 internal identifier: ps_avgpres_adrenalglandns13_u133p2) and Trunk Neural Crest (Etchevers) (92), NB datasets; Lastowska (93), Delattre (94), and Versteeg (95). For expression analysis of U/FAS and MYCN-regulated genes (*Figure 5d*): MYCN inhibition by the JQ-1 bromodomain inhibitor in BE(2)C NB cells (96).

Additional datasets from other sources: For analysis of U/FAS gene expression changes in response to TSmiR modulation (*Figure 4b*): Doxycycline induced *miR-22* expression in RD18 rhabdomyosarcoma cells (97), *miR-22* transfection in 293T human embryonic kidney cells (98), *LIN28A* knockdown in H9 human embryonic stem cells, which causes inhibition of LIN28A resulted in a significant increase in *let-7* family members (99,100), *let-7* transfection in human dermal lymphatic endothelial cells (public on GEO on Feb 23, 2023; not yet published), and PEO1 *miR-124* transfection of PEO1 ovarian cancer cells (101). For gene expression analysis of U/FAS- and MYC/N-regulated genes (*Figure 5c*): tetracycline “off” *MYC* induction in P493-6 human EBV-transformed B lymphocytes (102). For the adrenal gland and tibial nerve sample analysis (*Supplemental Figure S3*): data was obtained from the r2 database using GTEx Portal data, (v8, protein coding), dbGaP accession number phs000424.vN.pN. Adrenal gland and tibial nerve samples from ages 20-49 were used in our analysis. Additional details for GEO accession datasets used for secondary analyses are shown in *Supplemental Table S2*.

## Abbreviations

ALA: alpha-linolenic acid (18:3ω3)
AR: omega-6 arachidonic acid (20:4ω6)
DH: omega-3 docosahexaenoic acid (22:6ω3)
DTr: omega-9 docosatrienoic acid (22:3ω9)
EFAD: essential fatty acid deficiency
F: fatty acid(s)
FAS: *de novo* fatty acid synthesis (pathway)
FAS-core: genes, (*FASN, ELOVL6, SCD, FADS2, FADS1*)
FPKM: fragments per kilobase of exon per million mapped fragments
HUF: highly unsaturated fatty acids; 3 or more double bonds
L: linoleic acid (18:2ω6)
Mead: Mead acid (20:3ω9
NB: neuroblastoma
NHF: normal human fibroblasts
O: omega-9 oleic acid (18:1ω9)
TSmiRs: tumor suppressor microRNAs
UFAS: unsaturated fatty acid synthesis (pathways)
U/FAS: unsaturated and *de novo* fatty acid synthesis (pathways)
ω3: omega-3
ω6: omega-6
ω9: omega-9

## Acknowledgements

This study was supported by the Cancer Prevention and Research Institute of Texas; Grant RR180034 (PI: J.T.P).

