## Supplemental Material for "Unsaturated fatty acid synthesis is associated with poor prognosis and differentially regulated by *MYCN* and tumor suppressor microRNAs in neuroblastoma"

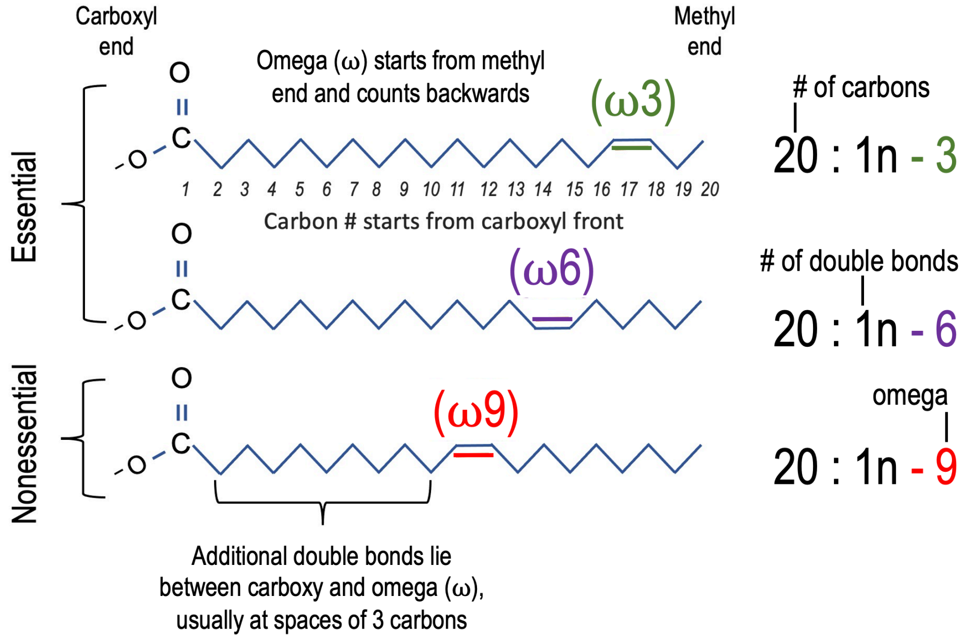


**Supplemental Figure S1. Unsaturated fatty acid structure.** Schematic of ω3, ω6, and ω9 fatty acid structures showing that carbon chains are numbered starting from the carboxy-terminus and that ω-double bonds are defined by distance from the terminal methyl group of the opposite end of the carbon chain. Nomenclature starts with number of carbons in the chain followed by a colon, then the total number "n" of double bonds in the chain, followed by a dashed number defining the ω class of the fatty acid.

| Structure | Common name | DB location, (Δ) | Component |
| --- | --- | --- | --- |
| **Saturated** | | | |
| SFA | myristic (MA) | — | 14:0 |
| SFA | palmitic (PA) | — | 16:0 |
| SFA | margaric | — | 17:0 |
| SFA | stearic (SA) | — | 18:0 |
| SFA | arachidic | — | 20:0 |
| SFA | behenic | — | 22:0 |
| SFA | lignoceric | — | 24:0 |
| **Omega-9 (ω9)** | | | |
| MUFA | cis-hexadecenoic | Δ7 | 16:1ω9 |
| MUFA | oleic (OA) | Δ9 | 18:1ω9 |
| MUFA | gondoic | Δ11 | 20:1ω9 |
| PUFA | 8,11-eicosadienoic | Δ8,11 | 20:2ω9 |
| HUFA | 5,8,11-eicosatrienoic (Mead) | Δ5,8,11 | 20:3ω9 |
| MUFA | erucic | Δ13 | 22:1ω9 |
| HUFA | 7,10,13-docosatrienoic (DTrA) | Δ7,10,13 | 22:3ω9 |
| MUFA | nervonic | Δ15 | 24:1ω9 |
| **Omega-7 (ω7)** | | | |
| MUFA | palmitoleic | Δ9 | 16:1ω7 |
| MUFA | asclepic | Δ11 | 18:1ω7 |
| **Omega-6 (ω6)** | | | |
| PUFA | linoleic (LA) | Δ9,12 | 18:2ω6 |
| HUFA | gamma-linolenic (GLA) | Δ6,9,12 | 18:3ω6 |
| PUFA | 11,14-eicosadienoic | Δ11,14 | 20:2ω6 |
| HUFA | sciadonic | Δ5,11,14 | 20:3ω6 |
| HUFA | dihomo-gamma-linolenic (DGLA) | Δ8,11,14 | 20:3ω6 |
| HUFA | arachidonic (ARA) | Δ5,8,11,14 | 20:4ω6 |
| HUFA | adrenic (AdrA) | Δ7,10,13,16 | 22:4ω6 |
| HUFA | osbond | Δ4,7,10,13,16 | 22:5ω6 |
| **Omega-3 (ω3)** | | | |
| HUFA | alpha-linolenic (ALA) | Δ9,12,15 | 18:3ω3 |
| HUFA | eicosapentaenoic (EPA) | Δ5,8,11,14,17 | 20:5ω3 |
| HUFA | docosapentaenoic (DPA) | Δ7,10,13,16,19 | 22:5ω3 |
| HUFA | docosahexaenoic (DHA) | Δ4,7,10,13,16,19 | 22:6ω3 |

**Supplemental Table S1. List of fatty acids analyzed in this study.** Table showing structure class (SFA = saturated fatty acid; MUFA = monounsaturated fatty acid; PUFA = fatty acids with two double bonds (double unsaturated); HUFA = fatty acids with three or more double bonds), common names and abbreviations, location of double bonds (location numbered from carboxy-terminus), and the component nomenclature. The “ω9” and “n-9” notations are interchangeable nomenclatures: e.g., 20:3ω9 = 20:3n-9 = Mead acid.

| Sample Name | Cell Type | Cell Line Designation | Sex | MYCN Status | Therapy Phase | Date Established | NB Cell Relationship | Research Resource Identifier (RRID) |
|---|---|---|---|---|---|---|---|---|
| NHF | Normal Human Fibroblast | BJ | male | — | — | 2000 | — | CVCL_3653 |
| BE1 | Neuroblastoma | SK-N-BE(1) | male | amplified | Dx | 1972 | paired | CVCL_9898 |
| BE2 | Neuroblastoma | SK-N-BE(2) | male | amplified | PD | 1972 | | CVCL_0528 |
| C15 | Neuroblastoma | CHLA-15 | female | non-amplified | Dx | 1988 | paired | CVCL_6594 |
| C20 | Neuroblastoma | CHLA-20 | female | non-amplified | PD | 1988 | | CVCL_6602 |
| KAN | Neuroblastoma | SMS-KAN | female | amplified | Dx | 1978 | paired | CVCL_7131 |
| KANR | Neuroblastoma | SMS-KANR | female | amplified | PD | 1978 | | CVCL_7132 |
| KCN | Neuroblastoma | SMS-KCN | male | amplified | Dx | 1979 | paired | CVCL_7133 |
| KCNR | Neuroblastoma | SMS-KCNR | male | amplified | PD | 1979 | | CVCL_7134 |

**Supplemental Table S2. Human cell lines used in this study.** All cell lines were grown in RPMI media supplemented with 10% fetal bovine serum. Sample name is used for abbreviating graphs and corresponds to a unique cell line designation. NB cells were derived from tumors of high-risk patients at dates provided and are classified by sex, *MYCN* status, and therapy phase (Dx = at diagnosis; PD = during progressive disease). Paired NB cell sets were isolated from a single patient at diagnosis and during progressive disease. All cell lines were tested for mycoplasma before and during use.

| Gene | Gene ID | Ensembl ID | Ensembl length (nt) | RefSeq ID | RefSeq length (nt) |
|---|---|---|---|---|---|
| ACACA | 31 | ENST00000353139.5 | 2349 | NM_198834.3 | 2324 |
| FASN | 2194 | ENST00000306749.2 | 2245 | NM_004104.5 | 805 |
| ELOVL6 | 79071 | ENST00000394607.3 | 5492 | NM_024090.3 | 5492 |
| SCD | 6319 | ENST00000370355.2 | 3901 | NM_005063.5 | 3893 |
| FADS2 | 9415 | ENST00000257261.6 | 2890 | NM_004265.4 | 1665 |
| ELOVL5 | 60481 | ENST00000370918.4 | 1741 | NM_021814.5 | 1723 |
| FADS1 | 3992 | ENST00000350997.7 | 5184 | NM_013402.7 | 2784 |
| ELOVL2 | 54898 | ENST00000354666.3 | **3027** | NM_0177770.4 | 3022 |
| MYCN | 4613 | ENST00000281043.3 | 917 | NM_001293228.2 | 907 |
| MYC | 4609 | ENST00000377970.2 | 1997 | NM_002467.6 | 1993 |
| MYCL | 4610 | ENST00000397332.2 | 1948 | NM_001033081.3 | 1944 |
| SDHA | 6389 | ENST00000264932.6 | 1269 | NM_004168.4 | 662 |
| ACTB | 60 | ENST00000331789.5 | 613 | NM_001101.5 | 600 |
| GAPDH | 2597 | ENST00000229239.5 | 224 | NM_002046.7 | 201 |
| DLK1 | 8788 | ENST00000341267.4 | 3326 | NM_003836.7 | 3328 |
| ITGAV | 3685 | ENST00000261023.3 | 3610 | NM_002210.5 | 3609 |
| PTEN | 5728 | ENST00000371953.3 | 6458 | NM_000314.8 | 6458 |

**Supplemental Table S3. Table of human genes used to identify potential 3’ UTR miRNA binding.** For each gene, gene symbol and ID are listed. The Ensemble ID and 3’UTR length are used to determine miRNA binding sites by TargetScan. Listed miRNA sites in *Table 1* were modified by limiting those sites to the RefSeq ID sequence and length, which are often shorter than the Ensembl sequences.

| Structure | Common name | Δ (DB location) | Component | NHF | BE1 | BE2 | C15 | C20 | KAN | KANR | KCN | KCNR |
|---|---|---|---|---|---|---|---|---|---|---|---|---|
| **Saturated** | | | | | | | | | | | | |
| SFA | myristic (MA) | — | 14:0 | 1.74 ± 0.10 | 1.49 ± 0.01 | 1.96 ± 0.05 | 2.32 ± 0.08 | 1.87 ± 0.05 | 2.33 ± 0.25 | 2.18 ± 0.31 | 3.29 ± 0.07 | 2.83 ± 0.01 |
| SFA | palmitic (PA) | — | 16:0 | 20.90 ± 0.37 | 19.12 ± 0.08 | 20.63 ± 0.06 | 24.22 ± 0.21 | 25.78 ± 0.07 | 26.74 ± 0.08 | 28.22 ± 0.90 | 23.09 ± 0.15 | 24.51 ± 0.25 |
| SFA | margaric | — | 17:0 | 0.75 ± 0.00 | 0.11 ± 0.02 | 0.14 ± 0.01 | 0.07 ± 0.01 | 0.14 ± 0.01 | 0.07 ± 0.03 | 0.19 ± 0.05 | 0.11 ± 0.04 | 0.17 ± 0.03 |
| SFA | stearic (SA) | — | 18:0 | 15.97 ± 0.35 | 16.96 ± 0.27 | 12.56 ± 0.11 | 12.37 ± 0.07 | 11.31 ± 0.04 | 10.52 ± 0.15 | 10.32 ± 0.49 | 14.02 ± 0.02 | 13.30 ± 0.01 |
| SFA | arachidic | — | 20:0 | 0.13 ± 0.00 | 0.12 ± 0.00 | 0.15 ± 0.02 | 0.16 ± 0.07 | 0.13 ± 0.02 | 0.20 ± 0.06 | 0.27 ± 0.08 | 0.09 ± 0.02 | 0.09 ± 0.02 |
| SFA | behenic | — | 22:0 | 0.45 ± 0.02 | 0.03 ± 0.02 | 0.15 ± 0.05 | 0.11 ± 0.04 | 0.27 ± 0.02 | 0.14 ± 0.00 | 0.07 ± 0.01 | 0.02 ± 0.00 | 0.09 ± 0.05 |
| SFA | lignoceric | — | 24:0 | 1.05 ± 0.01 | 0.04 ± 0.02 | 0.12 ± 0.00 | 0.17 ± 0.02 | 0.16 ± 0.06 | 0.07 ± 0.01 | 0.07 ± 0.01 | 0.06 ± 0.04 | 0.02 ± 0.00 |
| **Omega-7 (ω7)** | | | | | | | | | | | | |
| MUFA | palmitoleic | Δ9 | 16:1ω7 | 1.96 ± 0.04 | 0.59 ± 0.03 | 2.63 ± 0.01 | 2.43 ± 0.10 | 1.73 ± 0.25 | 1.79 ± 0.09 | 2.03 ± 0.02 | 1.53 ± 0.01 | 2.15 ± 0.12 |
| MUFA | asclepic | Δ11 | 18:1ω7 | 6.82 ± 0.01 | 7.08 ± 0.12 | 9.22 ± 0.08 | 7.54 ± 0.04 | 7.56 ± 0.03 | 8.91 ± 0.03 | 9.31 ± 0.27 | 8.56 ± 0.00 | 7.63 ± 0.10 |
| **Omega-9 (ω9)** | | | | | | | | | | | | |
| MUFA | cis-hexadecenoic | Δ7 | 16:1ω9 | 2.37 ± 0.05 | 3.57 ± 0.00 | 3.75 ± 0.02 | 6.26 ± 0.02 | 5.40 ± 0.17 | 5.32 ± 0.01 | 4.77 ± 0.19 | 4.64 ± 0.02 | 4.83 ± 0.01 |
| MUFA | oleic (OA) | Δ9 | 18:1ω9 | 20.98 ± 0.15 | 29.18 ± 0.27 | 32.42 ± 0.10 | 26.33 ± 0.08 | 24.31 ± 0.06 | 22.34 ± 0.05 | 22.03 ± 0.41 | 21.28 ± 0.19 | 26.24 ± 0.16 |
| MUFA | gondoic | Δ11 | 20:1ω9 | 0.28 ± 0.03 | 1.35 ± 0.00 | 1.58 ± 0.02 | 1.39 ± 0.06 | 1.21 ± 0.09 | 1.45 ± 0.04 | 0.88 ± 0.07 | 1.12 ± 0.02 | 0.73 ± 0.03 |
| PUFA | 8,11-eicosadienoic | Δ8,11 | 20:2ω9 | 0.23 ± 0.03 | 0.11 ± 0.02 | 0.36 ± 0.03 | 0.85 ± 0.03 | 0.60 ± 0.04 | 0.12 ± 0.07 | 0.08 ± 0.00 | 0.24 ± 0.04 | 0.32 ± 0.06 |
| HUFA | 5,8,11-eicosatrienoic (Mead) | Δ5,8,11 | 20:3ω9 | 0.17 ± 0.01 | 5.36 ± 0.22 | 3.80 ± 0.10 | 5.23 ± 0.04 | 4.72 ± 0.02 | 4.91 ± 0.00 | 2.15 ± 0.44 | 3.93 ± 0.08 | 2.54 ± 0.04 |
| MUFA | erucic | Δ13 | 22:1ω9 | 0.12 ± 0.03 | 0.07 ± 0.02 | 0.18 ± 0.01 | 0.20 ± 0.00 | 0.21 ± 0.05 | 0.10 ± 0.01 | 0.13 ± 0.01 | 0.05 ± 0.02 | 0.18 ± 0.05 |
| HUFA | 7,10,13-docosatrienoic (DTrA) | Δ7,10,13 | 22:3ω9 | 0.14 ± 0.01 | 5.96 ± 0.16 | 3.76 ± 0.08 | 3.76 ± 0.02 | 4.05 ± 0.01 | 5.75 ± 0.23 | 2.75 ± 0.43 | 5.42 ± 0.09 | 2.13 ± 0.07 |
| MUFA | nervonic | Δ15 | 24:1ω9 | 1.61 ± 0.01 | 0.10 ± 0.08 | 0.68 ± 0.03 | 0.59 ± 0.02 | 0.85 ± 0.02 | 0.31 ± 0.04 | 0.21 ± 0.03 | 0.12 ± 0.04 | 0.24 ± 0.02 |
| **Omega-6 (ω6)** | | | | | | | | | | | | |
| PUFA | linoleic (LA) | Δ9,12 | 18:2ω6 | 2.11 ± 0.05 | 0.03 ± 0.00 | 0.39 ± 0.44 | 0.13 ± 0.01 | 0.23 ± 0.00 | 0.32 ± 0.25 | 0.13 ± 0.07 | 0.08 ± 0.03 | 0.07 ± 0.00 |
| HUFA | gamma-linolenic (GLA) | Δ6,9,12 | 18:3ω6 | 0.30 ± 0.13 | 0.00 ± 0.00 | 0.00 ± 0.00 | 0.00 ± 0.00 | 0.00 ± 0.00 | 0.00 ± 0.00 | 0.00 ± 0.00 | 0.00 ± 0.00 | 0.00 ± 0.00 |
| PUFA | 11,14-eicosadienoic | Δ11,14 | 20:2ω6 | 0.09 ± 0.02 | 0.20 ± 0.06 | 0.69 ± 0.06 | 0.74 ± 0.02 | 0.54 ± 0.02 | 0.51 ± 0.02 | 0.18 ± 0.09 | 0.31 ± 0.02 | 0.07 ± 0.02 |
| HUFA | sciadonic | Δ5,11,14 | 20:3ω6 | 0.00 ± 0.00 | 0.00 ± 0.00 | 0.00 ± 0.00 | 0.00 ± 0.00 | 0.00 ± 0.00 | 0.00 ± 0.00 | 0.00 ± 0.00 | 0.00 ± 0.00 | 0.00 ± 0.00 |
| HUFA | dihomo-gamma-linolenic (DGLA) | Δ8,11,14 | 20:3ω6 | 1.90 ± 0.05 | 0.02 ± 0.02 | 0.11 ± 0.02 | 0.30 ± 0.04 | 0.47 ± 0.01 | 0.10 ± 0.05 | 0.18 ± 0.06 | 0.09 ± 0.01 | 0.12 ± 0.05 |
| HUFA | arachidonic (ARA) | Δ5,8,11,14 | 20:4ω6 | 10.28 ± 0.44 | 3.62 ± 0.02 | 2.35 ± 0.01 | 2.72 ± 0.02 | 4.67 ± 0.02 | 3.90 ± 0.01 | 7.64 ± 0.58 | 5.04 ± 0.06 | 5.85 ± 0.02 |
| HUFA | adrenic AdrA) | Δ7,10,13,16 | 22:4ω6 | 3.25 ± 0.07 | 2.13 ± 0.12 | 0.97 ± 0.11 | 0.79 ± 0.04 | 1.22 ± 0.06 | 0.71 ± 0.02 | 2.03 ± 0.16 | 3.46 ± 0.05 | 2.59 ± 0.06 |
| HUFA | osbond | Δ4,7,10,13,16 | 22:5ω6 | 0.83 ± 0.03 | 0.51 ± 0.08 | 0.09 ± 0.01 | 0.20 ± 0.03 | 0.59 ± 0.03 | 1.55 ± 0.07 | 1.71 ± 0.18 | 0.34 ± 0.02 | 0.42 ± 0.04 |
| **Omega-3 (ω3)** | | | | | | | | | | | | |
| HUFA | alpha-linolenic (ALA) | Δ9,12,15 | 18:3ω3 | 0.36 ± 0.06 | 0.00 ± 0.00 | 0.00 ± 0.00 | 0.00 ± 0.00 | 0.00 ± 0.00 | 0.00 ± 0.00 | 0.00 ± 0.00 | 0.00 ± 0.00 | 0.00 ± 0.00 |
| HUFA | eicosapentaenoic (EPA) | Δ5,8,11,14,17 | 20:5ω3 | 0.07 ± 0.05 | 0.00 ± 0.00 | 0.00 ± 0.00 | 0.00 ± 0.00 | 0.00 ± 0.00 | 0.00 ± 0.00 | 0.00 ± 0.00 | 0.00 ± 0.00 | 0.00 ± 0.00 |
| HUFA | docosapentaenoic (DPA) | Δ7,10,13,16,19 | 22:5ω3 | 2.51 ± 0.13 | 0.91 ± 0.03 | 0.59 ± 0.03 | 0.22 ± 0.01 | 0.46 ± 0.00 | 0.18 ± 0.08 | 0.46 ± 0.04 | 1.37 ± 0.04 | 1.22 ± 0.13 |
| HUFA | docosahexaenoic (DHA) | Δ4,7,10,13,16,19 | 22:6ω3 | 2.67 ± 0.19 | 1.35 ± 0.02 | 0.71 ± 0.01 | 0.90 ± 0.04 | 1.52 ± 0.05 | 1.66 ± 0.14 | 2.01 ± 0.14 | 1.73 ± 0.02 | 1.68 ± 0.12 |

**Supplemental Table S4. NB cell line lipid levels master table.** Fatty acids are ordered by structure and increasing component size. Numerical levels represent percent weight of a given fatty acid per overall weight (w/w, %) as determined by GC/MS (three biological replicates per cell line).

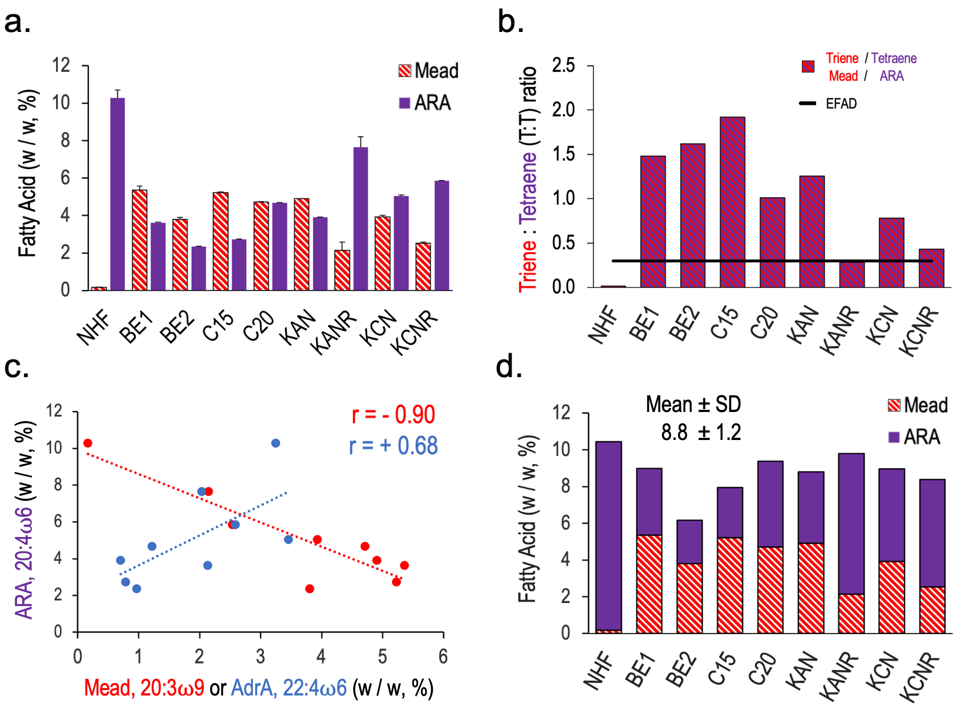


**Supplemental Figure S2. Comparison of ω9 Mead acid and ω6 ARA levels in human cell lines. a.** Comparison of Mead (red striped bars) and ARA (purple bars) levels in normal human fibroblasts (NHF) and human NB cell lines. **b.** Triene:Tetraene (T:T) ratios in human cell lines. Comparison of Mead (20:3ω9):ARA (20:4ω6**)** level ratios for NHF and human NB cell lines. Essential fatty acid deficiency (EFAD) is defined by a T:T ratio above 0.25 (denoted by black line). **c.** Correlation graphs of ARA levels to levels of Mead (red) or ARA-elongation product, AdrA (blue). **d.** Sum of Mead (red strips) and ARA (purple) levels across NHF and human NB cell lines. The mean and SD of all cell lines is 8.8±1.2 (w/w, %).

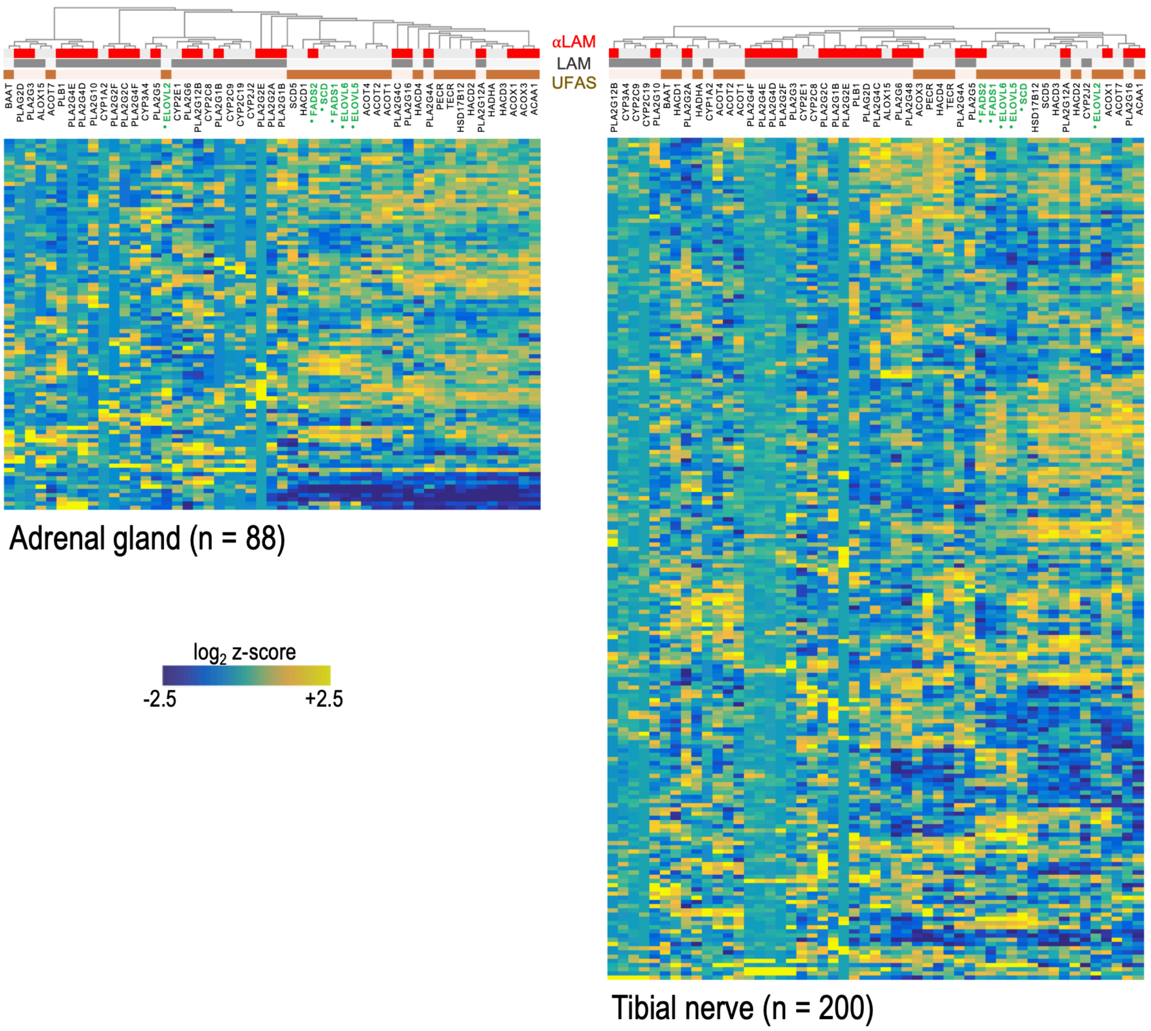


**Supplemental Figure S3. Expression patterns in normal human tissues by KEGG gene set analysis.** Heatmap of α-linolenic acid metabolism (αLAM), linoleic acid metabolism (LAM), and unsaturated fatty acid synthesis (UFAS) gene expression comparing adrenal gland cells and tibial nerve cells to other body tissues. U/FAS pathway genes from *figure 1b* are titled in green and contain asterisks (*). Genes associated with αLAM, LAM and U/FAS gene sets are identified by clustered red, grey, and brown bars, respectively. Expression data is presented as normalized values ($log_2$ z-scores). (*GTEx Portal v8, protein coding*)

a.

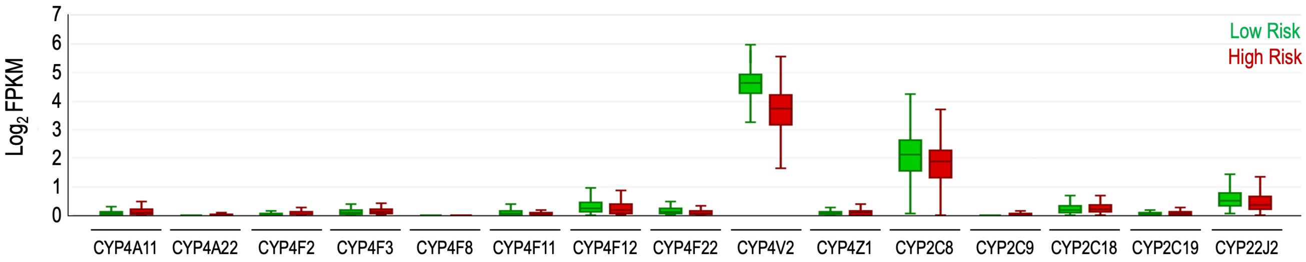


b.

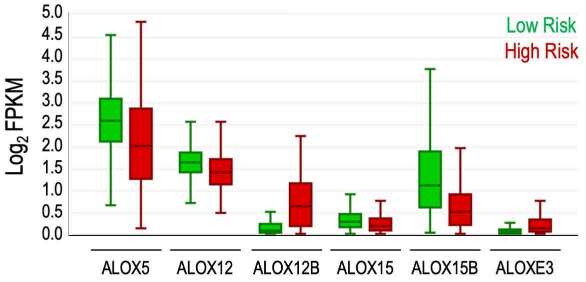


**Supplemental Figure S4. Expression level comparisons of CYP450 and LOX genes in low-risk and high-risk NB patients. a-b.** Expression levels of the *(a)* the fifteen human ω6 hydroxylating CYP450 and *(b)* the six human lipoxygenase (ALOX) genes in low-risk (green bars, n = 270) and high-risk (red bars, n = 171) NB. Gene expression is displayed for stage 1-4 NB patients by $\log_2$ FPKM values. (*GEO accession GSE62564*)

| Figure | Key Word(s) | GEO Accession | Authors | PMID | R2 Internal Identifier |
|---|---|---|---|---|---|
| 2a | NB | GSE62564 | SEQC (Su) | 25633159 | ps_avgpres_gse49710geo498_ag44kcwolf |
| 2b | NB | GSE62564 | SEQC (Su) | 25633159 | ps_avgpres_gse49710geo498_ag44kcwolf |
| 3a | NB | GSE62564 | SEQC (Su) | 25633159 | ps_avgpres_gse49710geo498_ag44kcwolf |
| 3b | NB | GSE62564 | SEQC (Su) | 25633159 | ps_avgpres_gse49710geo498_ag44kcwolf |
| 4b | *miR-22* | GSE83805 | Bersani | 27569217 | n/a |
| 4b | *miR-22* | GSE92599 | Inwood | 28987030 | n/a |
| 4b | *LIN28A (let-7)* | GSE53038 | Kim | 25479749 | n/a |
| 4b | *let-7* | GSE218501 | n/a | not yet published | n/a |
| 4b | *miR-124* | GSE67327 | Shields | 26655797 | n/a |
| 4c | NB | GSE62564 | SEQC (Su) | 25633159 | ps_avgpres_gse49710geo498_ag44kcwolf |
| 5a | Adrenal Gland | GSE3526 | Various | N/A | ps_avgpres_adrenalglandns13_u133p2 |
| 5a | Adrenal Gland | GSE7307 | Various | N/A | ps_avgpres_adrenalglandns13_u133p2 |
| 5a | Adrenal Gland | GSE8514 | Various | N/A | ps_avgpres_adrenalglandns13_u133p2 |
| 5a | Trunk Neural Crest | GSE14340 | Etchevers | 19666486 | ps_avgpres_gse14340geo5_u133p2 |
| 5a | NB | GSE16476 | Versteeg | 22367537 | ps_avgpres_nbadam88_u133p2 |
| 5a | NB | GSE13136 | Lastowska | 17533364 | ps_avgpres_nblastowska30_u133p2 |
| 5a | NB | GSE12460 | Delattre | 18553563 | ps_avgpres_nbdelatre64_u133p2 |
| 5b | NB | GSE62564 | SEQC (Su) | 25633159 | ps_avgpres_gse49710geo498_ag44kcwolf |
| 5c | *MYC* (induction) | GSE14302 | Chang | 19211792 | n/a |
| 5d | MYCN (JQ1) | GSE107707 | Iniguez | 30537514 | ps_avgpres_gse107707geo30_gpl18573b |
| 5e | NB | GSE62564 | SEQC (Su) | 25633159 | ps_avgpres_gse49710geo498_ag44kcwolf |
| 6b | NB | GSE62564 | SEQC (Su) | 25633159 | ps_avgpres_gse49710geo498_ag44kcwolf |
| 7 | NB | GSE62564 | SEQC (Su) | 25633159 | ps_avgpres_gse49710geo498_ag44kcwolf |
| Sup S4a | NB | GSE62564 | SEQC (Su) | 25633159 | ps_avgpres_gse49710geo498_ag44kcwolf |
| Sup S4b | NB | GSE62564 | SEQC (Su) | 25633159 | ps_avgpres_gse49710geo498_ag44kcwolf |

**Supplemental Table S5. Publicly available GSE data sets used in this study.** Data sets used for secondary analysis are organized by figure location and key words. Gene Expression Omnibus (GSE) accessions, study authors, PubMed Identifier (PMID), and R2 database internal identifiers are provided.
